# The role of mu opioid receptors on excitatory and inhibitory neurons in the rostral ventromedial medulla in neuropathic pain

**DOI:** 10.64898/2026.08.26.747135

**Authors:** Jamie C. Moffa, Amy Gao, Vani Kalyanaraman, Monique Heitmeier, Bryan Copits

## Abstract

Descending projections from the brain to the spinal cord can regulate painful stimulus processing and are modulated by endogenous and exogenous opioids. We investigated the role of mu opioid receptors (MORs) in GABAergic vs. glutamatergic neurons of the rostral ventral medulla (RVM) in a mouse model of chronic neuropathic pain. We found that activating glutamatergic and GABAergic neurons in the RVM both result in antinociception at baseline, but glutamatergic neurons *enhance* pain responses after nerve injury. We then interrogated the role of RVM MOR signaling on neuropathic pain by using CRISPR/Cas9 to delete MOR in glutamatergic or GABAergic RVM neurons. We found that MOR knockout in glutamatergic and GABAergic RVM neurons precipitates early neuropathic pain onset with no effect on chronic pain intensity. These results suggest that RVM MOR signaling modulates hypersensitivity in the early phase of injury, but chronic neuropathic pain is largely independent of mu opioid receptor signaling.

## Introduction

The body relies on a complex pain perception system to avoid serious injury. This system involves peripheral nociceptive neurons sending signals to the brain, and signals from the brain modifying information from those neurons. In chronic pain this system goes awry, resulting in persistent pain that impedes function. Chronic pain is a serious condition, affecting 50 million people in the United States annually [17]. However, the specific mechanisms underlying chronic pain, and how we can better treat it, are poorly understood.

The rostral ventromedial medulla (RVM) is a key hub in the descending pain pathway that receives input from upstream brain areas like the periaqueductal grey (PAG) and sends projections to the spinal cord [8,36,51], where it bi-directionally modulates pain signals [35]. Electrical or pharmacological RVM activation increases withdrawal thresholds in reflexive pain assays. Conversely, RVM inactivation has little effect at baseline, but is antinociceptive in chronic pain. [18,37,60,64]. The RVM densely expresses the mu opioid receptor (MOR) [46,59], which binds endogenous and exogenous opioids to modulate pain sensation. Opioid agonist injection into the RVM produces potent analgesia [1,5,19,20,41]; inactivation of the RVM or lesion of the dorsolateral funiculus prevents systemically administered opioids from achieving their full effect [32,61].

RVM neurons have been categorized by their responses to pain and opioid receptor agonists. “On-cells” are activated by noxious stimuli and inhibited by MOR agonist DAMGO; “Off-cells” are pain-inhibited and opioid-activated [2]. Neutral cells show no change in firing to either stimulus. On-, Off-, and neutral cells are all molecularly heterogenous: a majority of each functionally-defined cell type is GABAergic, with serotonergic neurons comprising a subset of neutral cells [68]. While useful, the on/off cell paradigm has limitations. First, most studies on RVM on- and off-cells were performed under anesthesia, which alters these populations’ stimulus responses [48,56,57]. Second, on-, off-, and neutral cells are difficult to target for modulation, and attempts to identify molecular markers of these populations have yielded conflicting results [25,42,55,69].

A different approach for understanding the RVM’s role in descending pain modulation instead focuses on GABAergic, glutamatergic, and serotonergic neurons. These neurons are easier to target using a combination of transgenic mouse lines and viral vectors. Activation and inhibition of each of these neuronal populations affects pain behaviors: GABAergic and glutamatergic neurons are broadly antinociceptive [21,29,55,71], while serotonergic neurons show complex effects on pain [3,11,31].

However, few studies have investigated the role of GABAergic and glutamatergic RVM neurons in neuropathic pain. Additionally, the role of MORs on these neurons has not been directly assessed. In this paper, we chemogenetically activate GABAergic and glutamatergic RVM neurons after inducing neuropathic pain. We then use CRISPR- Cas9 gene editing to selectively delete MORs in GABAergic or glutamatergic RVM neurons to assess the contribution of MOR signaling to the development of neuropathic pain. We uncover a switch from descending pain inhibition to facilitation in glutamatergic RVM neurons after neuropathic pain and show that MORs on GABAergic and glutamatergic RVM neurons play a role in early development, but not maintenance, of chronic neuropathic pain.

## Methods

### Model organisms

All procedures were conducted in accordance with National Institutes of Health guidelines and with approval from the Institutional Animal Care and Use Committee at Washington University in St. Louis. All mice were group housed and kept on a 12h light:12h dark cycle unless otherwise specified. For all experiments, male and female mice were used. For RVM *in situ* hybridization, C57Bl/6J (Jax #000664) mice were used. For chemogenetic activation experiments, Vgat-IRES-Cre (Slc32a1^tm2(cre)Lowl^, Jackson labs #028862) [66] or Vglut2-IRES-Cre (Slc17a6^tm2(cre)Lowl^; Jackson labs #016963) [66] mouse lines on a C57Bl/6 background were used to target GABAergic and glutamatergic neurons, respectively.

For CRISPR/Cas9 cell-type specific MOR knockout, the requisite mouse lines were generated by crossing homozygous H11-LSL-Cas9 knock-in mice (Igs2^tm1(CAG-Cas9*)/Mmw^; Jackson labs #027632) [13] to the following homozygous Cre lines: Vgat-IRES-Cre (Slc32a1^tm2(cre)Lowl^, Jackson labs #028862) [66] or Vglut2-IRES-Cre (Slc17a6^tm2(cre)Lowl^; Jackson labs #016963) [66].

### Viral constructs

AAV9-hSyn-DIO-mCherry, used as a control in chemogenetic experiments, was a gift from Bryan Roth (Addgene plasmid # 50459-AAV9; http://n2t.net/addgene:50459; RRID:Addgene_50459). AAV9-hSyn-DIO-hM3D(Gq)-mCherry, used in chemogenetic experiments, was a gift from Bryan Roth (Addgene plasmid # 44361-AAV9; http://n2t.net/addgene:44361; RRID:Addgene_44361) [44].

The EGFP-KASH sequence was derived from pAAV-FLEX-EGFP-KASH (gift from Larry Zweifel, Addgene #154373) and synthesized as a reverse transcribed G block from IDT technologies, with KpnI and NheI on the 5’ and 3’ ends respectively. We used pX552 DIO NLS-mRuby3 [50] as a vector for this G block. This vector was cut with KpnI and NheI and gel purified. pX552 vector was then ligated in the inverted orientation inside of the flanking loxP sites. The entire sequence was verified using forward primer 5’- CACCATCGACCCGAATTGCC-3’ and reverse primer 5’- GTGAGATCTGGACTAGAGGGTC-3’.

#### Designing MOR gRNA (gMOR)

gRNA candidates against the µ opioid receptor were designed as previously described [50]. Briefly, the first common exon among MOR isoforms in mouse was identified (in this case, exon 2). CRISPOR [16] was used to identify all possible gRNA candidates in MOR exon 2 that are upstream of the SpCas9 protospacer adjacent motif (PAM). Top candidates were selected based on their predicted efficiency and specificity and sent to the Genome Editing Stem Cell core at WashU to assess their editing efficiency in Neuro2A cells. The candidate with the best editing efficiency *in vitro* was selected, synthesized (along with a control replacing the last 3 nucleotides with ‘TTA’ to prevent binding near the SpCas9 PAM), and packaged into an empty U6-gRNA-EF1-DIO-eGFP-KASH vector (Addgene #260895) as previously described [38,39,49,50]. We used AAV-ITR sequencing (Azenta) to verify viral packaging sequences and expression cassettes. Active gMOR (Addgene #260893) and control TTA (Addgene #260894) vectors were packaged into AAV9 viruses, purified, concentrated, and stored at -80°C [39,50].

### Histology

#### RVM *in situ* hybridization

Adult male and female C57Bl/6 mice were deeply anesthetized with a ketamine/xylazine cocktail and perfused using 40 mL cold 1X phosphate buffered saline (PBS) and 40 mL cold 4% paraformaldehyde (PFA). After perfusion, brains were post-fixed at 4°C overnight in 4% PFA, then transferred to 30% sucrose solution and stored at 4°C overnight for cryoprotection. Tissue was then embedded in OCT and frozen on dry ice to prepare for cryostat sectioning. 20µm-thick coronal sections were taken of the RVM and stored in 1X PBS, then mounted on a positively-charged glass slide and allowed to dry completely.

Slides containing RVM sections ranging from 6.4mm posterior to Bregma to 5.6mm posterior to Bregma from each brain were prepared in triplicate to facilitate three co-labeling conditions. One set of slides were co-labeled for *Vgat* and *Oprm1* using probes from ACD bio, the second set co-labeled for *Vglut2* and *Oprm1*, and the third for *Tph2* and *Oprm1*. Amplified *Vgat*, *Vglut2*, or *Tph2* mRNA were stained using Opal 650 dye. In all conditions, amplified *Oprm1* mRNA was stained using Opal 570 dye. *In situ* hybridization (ISH) was performed according to the manual for the RNAScope Multiplex Fluorescent Reagent Kit v2 for PFA-fixed frozen sections. After completing ISH, a coverslip was applied using DAPI-containing mounting media for nuclear staining and slides were allowed to dry overnight.

#### RVM cFos Immunohistochemistry

Adult male and female *Vgat*-Cre mice underwent spared nerve injury (SNI) or sham surgery 10 days prior to perfusion and tissue harvesting. On day 10, mice were perfused and their brains post-fixed and cryoprotected as described above. 40µm coronal sections were prepared on the cryostat and stored in 1X PBS. For cFos immunostaining, RVM sections were first washed 3x for 10 minutes each in 1X PBS. Sections were then blocked and permeabilized in 5% normal donkey serum (NDS) and 0.5% TritonX for 1 hour at room temperature. Sections were transferred to 0.1% rabbit anti-phospho-cFos primary antibody (Cell Signaling #D82C12) in blocking buffer and incubated overnight at room temperature.

The next day, sections were washed 3x for 10 minutes each in 1X PBS before being transferred to secondary antibody solution containing 0.1% donkey anti-rabbit antibody conjugated to AF647 (Invitrogen #A31573) in blocking buffer. Sections were incubated in this solution for 2 hours at room temperature, then washed again 3x for 10 minutes each in 1X PBS. Sections were mounted on positively charged glass slides, a coverslip was applied using DAPI-containing mounting media for nuclear staining and slides were allowed to dry overnight.

#### Spinal Cord mCherry Immunohistochemistry

Adult male and female *Vgat-*Cre or *Vglut2*-Cre mice were injected with 100 nL AAV9- DIO-mCherry virus in the RVM 4 weeks before tissue harvesting and immunostaining. 4 weeks after virus injection, mice were perfused as described above, and their brains and spinal columns were post-fixed and cryoprotected as above. After cryoprotection, brains were mounted in OCT and 40µm RVM sections were taken and mounted to confirm virus injection coordinates.

Spinal columns were cleaned and a dorsal laminectomy performed to expose the spinal cord. ∼2cm of spinal cord tissue around the lumbar enlargement were dissected out and embedded in OCT. 40µm sections were taken from lumbar spinal cord levels L1-L5 and stored in 1X PBS. For mCherry immunostaining, the same procedure as above was followed, using 0.1% chicken anti-mCherry (Abcam # ab205402) in blocking buffer as the primary antibody, and 0.5% donkey anti-chicken conjugated to Cy3 (Jackson Laboratories # 703-165-155) plus 0.5% IB4 conjugated to AF 647 (Invitrogen #I32450) in blocking buffer as the secondary antibody. Tissues were mounted using DAPI mounting media and allowed to dry overnight.

### Imaging and analysis

#### RVM ISH Images and analysis

RVM sections co-labeled for *Vgat*, *Vglut2*, or *Tph2* and *Oprm1* were imaged on a Leica confocal microscope using a 40X oil immersion lens. DAPI was excited using the 405nm laser, Opal 570 dye (staining *Oprm1*) was excited using the 561nm laser, and Opal 650 dye (staining *Vgat*, *Vglut2*, or *Tph2*) was excited using the 638nm laser. Laser intensity, gain, and emission spectra were selected to minimize channel crosstalk while maximizing true signal, and values were kept consistent across all sections and conditions for quantification.

Cellpose software [63] was used to draw regions of interest (ROIs) around *Vgat*+, *Vglut2*+, or *Tph*2+ neurons, highlighting GABAergic, glutamatergic, and serotonergic RVM neurons, respectively. ROIs were imported into ImageJ and superimposed on the near-red *Oprm1* channel. The *Oprm1* images were background-subtracted and binarized (threshold intensity value of 30) such that pixel values above 30 were set to max (255) and pixel values below 30 were set to 0. The Analyze Puncta command was then used to count individual *Oprm1* puncta in each cell body ROI. Neurons with 1 or more *Oprm1* puncta were considered *Oprm1+*, and neurons with 0 puncta were considered *Oprm1* negative. The fraction of *Vgat*, *Vglut2*, and *Tph2* (lateral and medial) neurons positive for *Oprm1* were quantified for each animal and compared using a one-way ANOVA with follow-up t-tests with corrections for multiple comparisons.

Total *Vgat*+, *Vglut2*+, and *Tph2*+ neurons in each section were determined by adding the total neurons within the boundaries of the RVM for each section. Sections were then compared to Allen brain atlas to determine approximate rostral-caudal level relative to bregma. Since *Oprm1* mRNA staining does not reliably label cell bodies like the other 3 markers, total *Oprm1* puncta within the RVM per section was determined instead, using the thresholded images above.

#### RVM cFos imaging and analysis

RVM sections stained for phospho-cFos were imaged on a Keyence epifluorescence microscope. DAPI was imaged with blue dichroic filter, and AF647 (cFos) was imaged with a far-red dichroic filter. Stitched images were loaded into ImageJ, and ROIs were drawn around the RVM. Images were background subtracted and binarized at a pixel intensity value of 20 (pixels with a value <20 were set to 0 and pixels with a value >20 were set to maximum value). The Analyze Puncta function was used to count the number of cFos+ neurons within the defined ROI. Puncta count was normalized to the ROI area, giving the final value of cFos+ cells/mm^2^. The number of RVM cFos+ neurons between SNI and sham groups was compared using a nested t-test.

#### mCherry spinal cord imaging and analysis

Spinal cord sections stained for mCherry and IB4 were imaged on a Leica confocal microscope using a 20X air lens. DAPI was excited using the 405nm laser, mCherry was excited using the 561nm laser, and IB4-647 was excited using the 638nm laser. As above, laser intensity, gain, and emission spectra were selected to minimize crosstalk and autofluorescence and maximize true signal, and the same settings were used across all images in both Vgat-Cre and Vglut2-Cre sections to allow for appropriate quantification.

After imaging, stitched images of entire spinal cord sections were loaded into ImageJ. ROIs for dorsal horn laminae were determined as follows: **Laminae I-II (LI-II):** LI-II were defined as the outermost section of the dorsal horn to the inner blade of LII, using IB4 fluorescence as a guide to the inner demarcation line. **Laminae III-VI:** LIII-VI were defined from the region just ventral to the inner blade of LII (as determined by IB4 staining) to approximately the midline of the spinal cord, bisected horizontally, excluding the area around the central canal. **Lamina X:** LX was defined as the approximately circular region surrounding the central canal, with its radius defined as the distance from the central canal to the medial white matter dorsal to the central canal.

These ROIs were superimposed on the mCherry channel. After background subtraction, the raw integrated density (RID), defined as the sum of the intensity values of all pixels in the ROI, was calculated and normalized to the area of the region, giving the final measurement of RID/µm^2^. The mCherry channel was then binarized, with pixel values below 20 set to 0 and values above 20 set to the maximum pixel value (255). The %Area of mCherry fibers in each ROI was then calculated. Both measurements (RID/ µm^2^ and mCherry %Area) were compared for *Vgat*-Cre and *Vglut2*-Cre animals across laminae using a repeated measures one-way ANOVA, followed by post-hoc t-tests with correction for multiple comparisons.

#### Injection site confirmation

RVM sections for all virus injection experiments were imaged on a Leica confocal microscope using a 20X objective. In all sections, DAPI was excited using the 405nm laser. For mice injected with an eGFP expression vector, eGFP was excited using the 488nm laser. For mice injected with an mCherry vector, mCherry was excited using the 561nm laser. As above, laser intensity, gain, and emission spectra were selected to minimize cross-talk and autofluorescence and maximize true signal. Mice were excluded from all subsequent analysis if they exhibited no or minimal (<50 total neurons) eGFP/mCherry expression across the rostro-caudal extent of the RVM.

### Surgical procedures

#### Intracranial virus injections

Animals were prepared for stereotaxic surgery as described previously [50]. Briefly, mice were initially anesthetized in an induction chamber with 5% isoflurane (iso) inhaled anesthesia. They were then transferred to a heated pad on the stereotax, where iso anesthesia was maintained at 1.5-2% for the duration of surgery. Mice received 0.05mg/kg subcutaneous buprenorphineSR (sustained release) as pre-operative analgesia. After stabilizing thena skull, and the fur on the surface of the scalp was removed using Nair. Exposed skin was disinfected using Iodine solution followed by 70% ethanol. A midline incision in the scalp was made using sterile forceps and spring scissors. The skull was then balanced such that Bregma and lambda were level, and the left and right sides of the skull on either side of the midline suture were level. A sterile micro-drill was used to make a hole in the skull above the injection coordinates, and 100nL of virus was injected using a 1µl Hamilton syringe affixed to the stereotax. RVM injection coordinates (relative to Bregma) were: Anterior/Posterior: -6mm, Medial/Lateral: 0mm, Dorsal/Ventral: -6mm. Virus was injected at a rate of 50nL/minute, and the syringe was allowed to dwell in the injection site for 10 additional minutes to allow virus to spread before withdrawal. After virus injection, the scalp incision was sutured using sterile 6-0 nylon sutures, and the mouse was allowed to recover in a heated recovery chamber.

#### Spared Nerve Injury

For spared nerve injury (SNI) surgery, mice were initially anesthetized in an induction chamber with 5% iso, then transferred to a nose cone on a heating pad where anesthesia was maintained at 1.5-2% iso for the remainder of the procedure. Hair on the left hindleg was removed using Nair, and the skin was disinfected using Iodine solution and 70% ethanol. An incision was made in the left hindleg, approximately 1-2 mm below the iliac crest, to expose the muscle underneath. Sterile forceps were used to dissect down through the muscle and expose the three branches of the sciatic nerve. For sham surgeries, the nerve was gently lifted using the forceps. For full SNI surgery, the tibial and common peroneal branches of the nerve were isolated from the sural nerve and ligated using sterile 6-0 silk sutures. The tibial and common peroneal nerves were then cut using sterile spring scissors, taking care to avoid transecting nearby blood vessels. The severed (or intact, in the case of sham surgery) nerve was then replaced. The skin incision was closed using 6-0 nylon sutures, and mice were allowed to recover in a heated recovery chamber.

#### Behavioral Assays and Analysis

For all behavioral assays, mice were habituated to the experimenter, testing room, and individual plexiglass testing chambers for 1 hour/day for 2 days prior to testing. During habituation and testing, animals were prevented from seeing each other by placement of opaque barriers between each testing chamber.

##### Von Frey

For assessment of mechanical withdrawal thresholds, the up/down von Frey assay was used [12]. Briefly, mice were placed individually in plexiglass testing chambers on a mesh surface. On the day of testing, mice were habituated to the chambers for 1 hour before the assay began. Both the left and right hindpaws were tested, and responses were recorded and analyzed to determine the average 50% withdrawal threshold across 3 trials/paw. For MOR knockout studies, results were compared between control and edited groups using a t-test with Welch’s correction for unequal variance. For chemogenetic activation studies, thresholds were compared between control and HM3D(q) infected groups 30 minutes after receiving either i.p. saline or 0.1mg/kg DCZ using a 2-way ANOVA with matching for the saline vs. DCZ condition. Individual groups were then compared using post-hoc t-tests with correction for multiple comparisons.

##### Hargreaves

For assessment of thermal withdrawal latencies, the Hargreaves assay was used [34]. Mice were placed in individual plexiglass testing chambers separated by opaque barriers on a Hargreaves apparatus that was warmed to 23°C. As with von Frey, they were allowed to habituate to the chambers for 1 hour before testing. Thermal stimulus intensity was set to 18, and the thermal stimulus was applied to the right and left hindpaw for 3-5 trials each paw. Withdrawal latency was recorded for each trial for each paw as the time to lifting, flinching, licking, or biting the stimulated paw. The stimulus was cut off at 20s if there was no response to avoid tissue damage. Hargreaves withdrawal latencies for control and edited groups were compared using a t- test with Welch’s correction for unequal variances.

##### Hot Plate

For assessment of thermal withdrawal latencies, the hot plate assay was used. Mice were allowed to habituate to the testing room for 1 hour before the first trial. Before the first trial, the hot plate was set to 55°C. For each of 3 trials, mice were placed on the center of the enclosed hot plate, and the time to first nocifensive response was recorded. Paw licking, flinching/shaking, biting, or lifting were considered responses. Mice were removed after response or after 30 seconds if they did not respond, to avoid tissue damage. Animals were also tested for 1 trial 30 minutes after receiving 10 mg/kg i.p. morphine. Average response latencies for edited and control groups were compared using a t-test with Welch’s correction for unequal variances. Response latencies before and after morphine administration were compared using a 2-way ANOVA with matching across baseline and morphine conditions. Post-hoc t-tests with correction for multiple comparisons were used to compare individual groups.

##### Brush stroke

The brush stroke assay was used to test dynamic allodynia before and after induction of neuropathic pain via SNI. Similar to von Frey, mice were placed in plexiglass testing chambers separated by opaque barriers on a mesh surface, and allowed to habituate for 1 hour before testing. A fine paintbrush was used to stimulate the lateral plantar surface of the left and right hindpaws, for 3 trials each paw. Responses were scored on a scale from 0-3, where 0 was defined as no response, moving away from the stimulus, or brief (<1s) paw lifting; 1 was defined as prolonged paw lifting >1s; 2 was defined as lifting the stimulated paw above the plane of the body; and 3 was defined as flinching, licking, or biting the stimulated paw. For MOR knockout experiments, data were analyzed using a 3-way ANOVA to compare edited vs. control groups, SNI vs. sham, and baseline vs. post-SNI time points, with matching by time point. For chemogenetic activation experiments, since all animals received SNI, data were analyzed using a 2-way ANOVA to compare HM3D(Gq) vs. mCherry groups and baseline vs. post-SNI time points, with matching by time point. For all experiments, post-hoc t-tests were used to compare individual groups, with correction for multiple comparisons.

##### Acetone

The acetone assay was used to test cold hyperalgesia before and after induction of neuropathic pain via SNI. Similar to von Frey, mice were placed in plexiglass testing chambers separated by opaque barriers on a mesh surface, and allowed to habituate for 1 hour before testing. A 10mL syringe was filled with acetone and used to apply a drop of acetone to the left and right hindpaws, for 1 trial each paw. Mice were recorded on video, and the number of licking and flinching bouts were quantified for each mouse after the fact. Baseline differences in acetone response between edited and control or HM3D(Gq) and mCherry groups were compared using a t-test with Welch’s correction for unequal variance. For MOR knockout experiments, data were analyzed using a 3-way ANOVA to compare edited vs. control groups, SNI vs. sham, and baseline vs. post-SNI time points, with matching by time point. For chemogenetic activation experiments, since all animals received SNI, data were analyzed using a 2-way ANOVA to compare HM3D(Gq) vs. mCherry groups and baseline vs. post-SNI time points, with matching by time point. For all experiments, post-hoc t-tests were used to compare individual groups, with correction for multiple comparisons.

### Data and code availability

All data and analysis codes are available from the Lead Contact. All DNA and viral constructs generated here are deposited on Addgene and are also available upon request from the Lead Contact.

## Results

### Chemogenetic activation of GABAergic RVM neurons is anti-nociceptive but does not alter hypersensitivity in a neuropathic pain model

GABAergic neurons in the RVM (RVM_GABA_) are broadly antinociceptive in the absence of chronic pain [29,55,71]. However, much less is known about their contribution to neuropathic pain phenotypes. To address this question, we injected an AAV virus containing Cre-dependent G_q_-coupled DREADDs (AAV9-DIO-hM3Dq-mCherry) or Cre-dependent mCherry control (**Figure 1A, left**) into the RVM of adult Vgat-IRES-Cre mice (**Figure 1A, right**). After 4 weeks, we performed baseline acetone, von Frey, and brush stroke assays to assess cold, mechanical, and dynamic response thresholds, respectively (**Figure 1B**). Each assay was performed after intraperitoneal (i.p.) administration of saline (Sal), then repeated after deschloroclozapine (DCZ; 0.1 mg/kg) injection to activate Gq-DREADDs in RVM_GABA_ neurons (**Figure 1C**). We found that chemogenetic activation of RVM_GABA_ neurons reduced the number of licking and flinching bouts in response to acetone stimulation (**Figure 1D**) and increased the mechanical withdrawal threshold (**Figure 1E**) in hM3Dq-injected, but not mCherry control mice. As mice do not exhibit nocifensive responses to brush stroke at baseline, these data are not shown.

**Figure 1.**
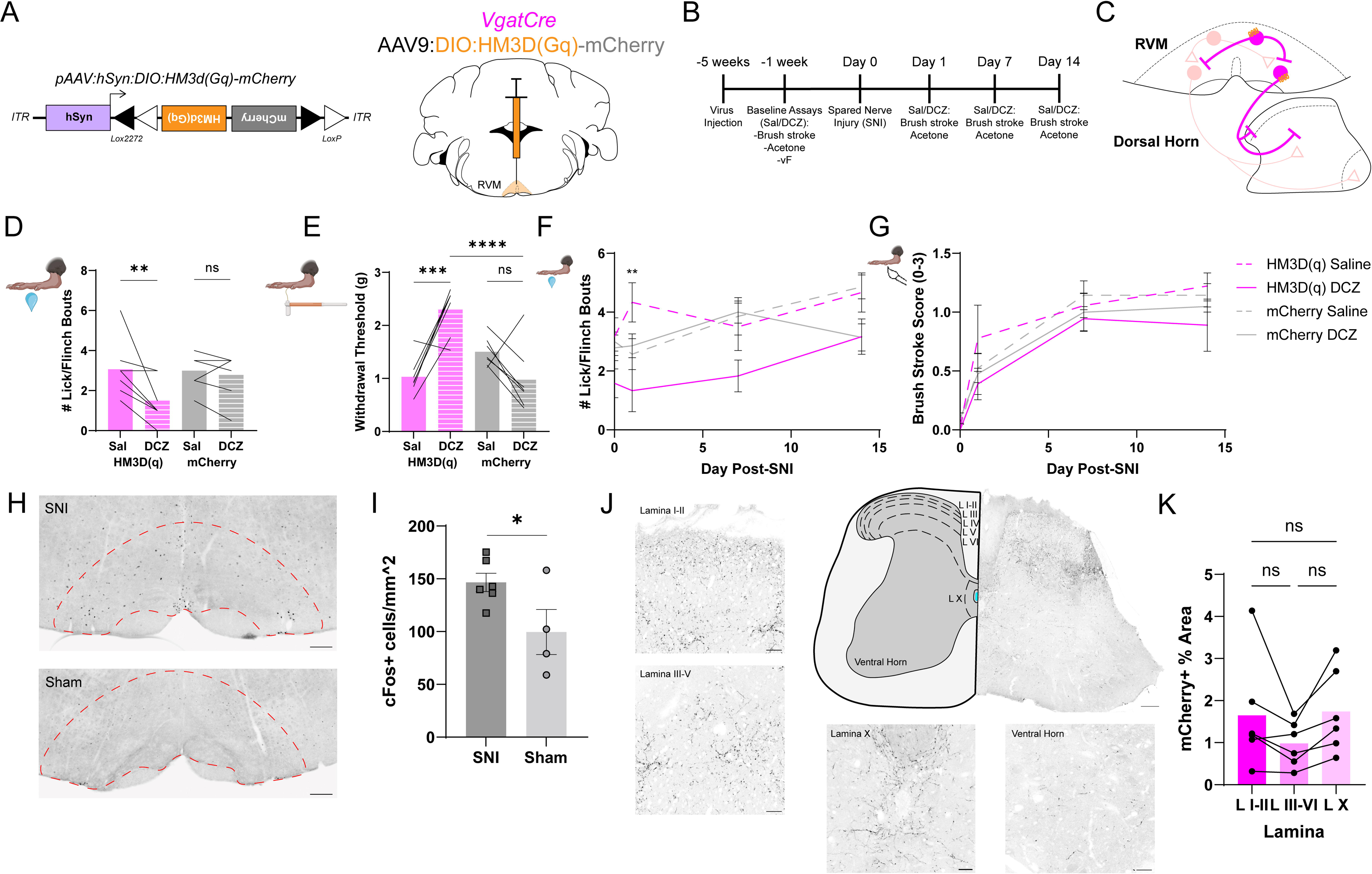
Chemogenetic activation of RVM_GABA_ neurons is antinociceptive at baseline with no effect after SNI. **A,** Diagram of AAV9 viral vector containing Cre- dependent, Gq-coupled DREADD HM3D(Gq) conjugated to mCherry (left); diagram of viral injection into the RVM of *Vgat-*Cre mice to target RVM_GABA_ neurons (right). **B,** Timeline of virus injection, behavioral assays, and spared nerve injury (SNI) procedure. **C,** Diagram of RVM_GABA_ neurons activated by deschloroclozapine (DCZ) injection in HM3D(Gq)-injected mice. **D,** Quantification of baseline number of licking/flinching bouts in response to acetone test in HM3D(Gq) (magenta) and mCherry control (gray) animals after injection of saline (sal; solid bars) or DCZ (striped bars). 2-way ANOVA with post- hoc t-tests corrected for multiple comparisons; **p<0.01. **E,** Quantification of baseline von Frey withdrawal threshold in HM3D(Gq) and mCherry control animals after injection of saline (sal) or DCZ. 2-way ANOVA with post-hoc t-tests corrected for multiple comparisons; ***p<0.001. **F,** Quantification of acetone assay licking/flinching bouts for HM3D(Gq) (magenta) and mCherry control (gray) mice after saline (dashed line) and DCZ (solid line) injection on days 1, 7, and 14 post-SNI. 3-way ANOVA with post-hoc t- tests corrected for multiple comparisons; **p<0.01 for HM3D(Gq) saline vs. DCZ on Day 1. **G,** Quantification of brush stroke assay score for HM3D(Gq) (magenta) and mCherry control (gray) mice after saline (dashed line) and DCZ (solid line) injection on days 1, 7, and 14 post-SNI. 3-way ANOVA with post-hoc t-tests corrected for multiple comparisons. **H,** Representative images of phospho-cFos immunostaining in the RVM of mice that underwent SNI (top) or sham surgery (bottom). Dashed red line indicates RVM. Scale bars=200µm. **I,** Quantification of the number of cFos+ cells/mm^2^ in the RVM of SNI mice vs. sham controls. Nested t-test, *p<0.05. **J,** Representative images of projection patterns of mCherry-expressing RVM_GABA_ neurons to the lumbar spinal cord. Main figure scale bar=100µm. From top left to bottom right, inset figures show enlarged images of RVM_GABA_ projection patterns in lamina I-II of the dorsal horn, lamina III-V of the dorsal horn, lamina X and the central canal of the spinal cord, and the ventral horn (scale bars=30µm). **K,** Quantification of the % Area of mCherry-expressing RVM_GABA_ projections in lamina I-II vs. lamina III-VI vs. lamina X of the dorsal horn of the lumbar spinal cord. Repeated measures one-way ANOVA. See **Supplemental Table 1** for detailed statistical analyses.

One week after baseline testing, we induced a neuropathic pain-like phenotype by performing spared nerve injury (SNI) surgery. We then repeated the acetone and brush stroke assays after saline and DCZ injection on days 1, 7, and 14 post-SNI. In the acetone assay, hM3Dq mice had significantly fewer licking and flinching bouts after DCZ administration (solid magenta line) compared to the control saline injection (dashed magenta line) on day 1 post-SNI (**Figure 1F**). Chemogenetic RVM_GABA_ activation did not have a significant effect on cold allodynia on days 7 and 14, nor did it affect dynamic allodynia measured by brush stroke response scores (**Figure 1G**). DCZ injection did not have a significant effect on either assay for mCherry controls (solid and dashed gray lines).

To confirm that SNI engages the RVM in descending pain modulation, we performed SNI or a control sham surgery on a separate cohort of mice. After 10 days to allow for maximum development of neuropathic pain, we stained RVM sections with an antibody to c-Fos, an immediate early gene and marker of neuronal activity (**Figure 1H**). We found a significant increase in cFos+ cells in SNI mice compared to sham controls (**Figure 1I**), suggesting that the neuropathic pain-like phenotype caused by SNI engages descending pain modulation networks within the RVM.

We next analyzed the distribution of RVM_GABA_ descending projections in the lumbar spinal cord from Vgat-Cre mice expressing cytosolic mCherry. In accordance with previous studies [27,30,53,71], we found spinally-projecting RVM_GABA_ neuronal processes in LI-II (**Figure 1J, left top**) and LIII-V (**Figure 1J, left bottom**). Additionally, we found processes in LX around the central canal (**Figure 1J, bottom left**), and throughout the ventral horn (**Figure 1J, bottom right**). The RVM_GABA_ fiber density in each level of the dorsal horn is quantified in **Figure 1K**, showing no significant differences in fiber density between LI-II, LIII-V, and LX.

### Chemogenetic activation of glutamatergic RVM neurons produces baseline antinociception, but exacerbates allodynia after nerve injury

To determine how glutamatergic neurons in the RVM (RVM_Glu_) influence baseline sensory thresholds and neuropathic pain, we expressed Cre-dependent Gq-coupled DREADDs (AAV9-DIO-hM3Dq-mCherry) or control virus (AAV9-DIO-mCherry) into the RVM of Vglut2-IRES-Cre mice (**Figure 2A**). After 4 weeks, we performed acetone, von Frey, and brush stroke assays after i.p. saline and DCZ (0.1 mg/kg) to activate Gq- DREADDs in RVM_Glu_ neurons (**Figure 2B**). Unlike RVM_GABA_ neurons, chemogenetic activation of RVM_Glu_ neurons had no effect on the number of licking and flinching bouts in response to acetone (**Figure 2C**). However, DCZ injection increased von Frey thresholds in hM3Dq-expressing mice, compared to control saline injection (**Figure 2D**). In both assays, DCZ had no effect on control mCherry-expressing mice.

**Figure 2.**
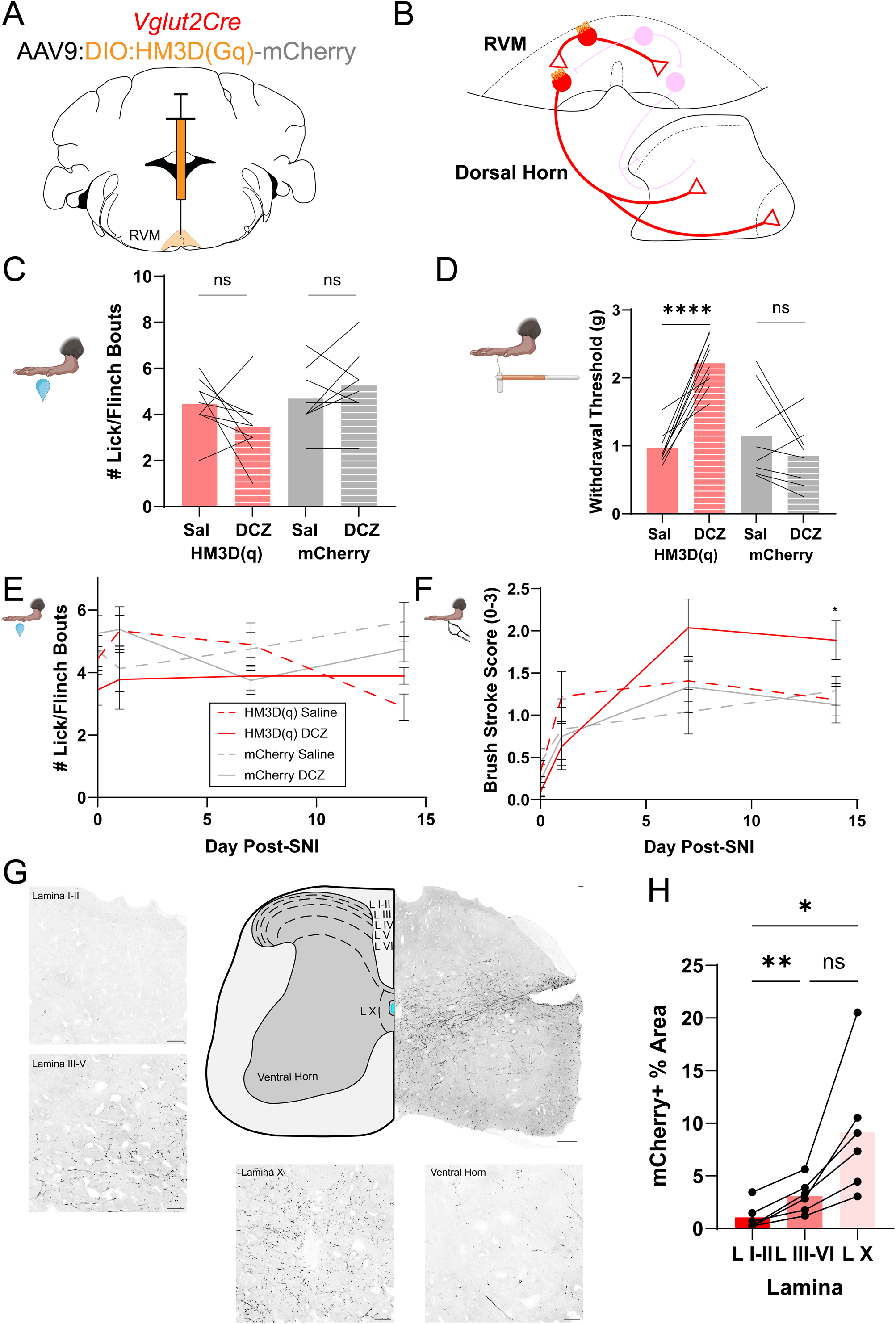
Chemogenetic activation of RVM_Glu_ neurons is antinociceptive at baseline but pro-nociceptive after SNI. **A,** Diagram of AAV9:DIO:HM3D(Gq)-mCherry viral injection into the RVM of *Vglut2-*Cre mice to target RVM_Glu_ neurons. **B,** Diagram of RVM_Glu_ neurons activated by deschloroclozapine (DCZ) injection in HM3D(Gq)-injected mice. **C,** Quantification of baseline number of licking/flinching bouts in response to the acetone test in HM3D(Gq) (red) and mCherry control (gray) animals after injection of saline (sal; solid bars) or DCZ (striped bars). 2-way ANOVA with post-hoc t-tests corrected for multiple comparisons. **D,** Quantification of baseline von Frey withdrawal threshold in HM3D(Gq) and mCherry control animals after injection of saline (sal) or DCZ. 2-way ANOVA with post-hoc t-tests corrected for multiple comparisons, ****p<0.0001. **E,** Quantification of acetone assay licking/flinching bouts for HM3D(Gq) (red) and mCherry control (gray) mice after saline (dashed line) and DCZ (solid line) injection on days 1, 7, and 14 post-SNI. 3-way ANOVA with post-hoc t-tests corrected for multiple comparisons. **F,** Quantification of brush stroke assay score for HM3D(Gq) (red) and mCherry control (gray) mice after saline (dashed line) and DCZ (solid line) injection on days 1, 7, and 14 post-SNI. 3-way ANOVA with post-hoc t-tests corrected for multiple comparisons; *p<0.05 for HM3D(Gq) saline vs. DCZ on Day 14. **G,** Representative images of projection patterns of mCherry-expressing RVM_Glu_ neurons to the lumbar spinal cord. Main figure scale bar=100µm. From top left to bottom right, inset figures show enlarged images of RVM_Glu_ projection patterns in lamina I-II of the dorsal horn, lamina III-V of the dorsal horn, lamina X and the central canal of the spinal cord, and the ventral horn (scale bars=30µm). **H,** Quantification of the %Area of mCherry- expressing RVM_Glu_ projections in lamina I-II vs. lamina III-VI vs. lamina X of the dorsal horn of the lumbar spinal cord. Repeated measures one-way ANOVA with post-hoc t- tests with correction for multiple comparisons; *p<0.05, **p<0.01. See **Supplemental Table 2** for detailed statistical analyses.

We performed SNI surgery 1 week later, and repeated the acetone and brush stroke assays with saline and DCZ injection on days 1, 7, and 14 post-SNI. RVM_Glu_ neuronal activation did not significantly affect acetone responses at any time point post-SNI (**Figure 2E**). In the brush stroke assay, DCZ injection in the hM3Dq group (**Figure 2F, solid red line**) had no effect on allodynia on days 1 or 7 post-SNI compared to saline alone (**Figure 2F, dashed red line**). However, on day 14 post-SNI, DCZ injection resulted in an *increased* score in response to brush stroke compared to controls, indicating that driving RVM_Glu_ activity exacerbates tactile allodynia following SNI. DCZ injection had no effect on mCherry controls in either assay (solid and dashed gray lines).

Compared to RVM_GABA_ neurons, much less is known about the descending projections from glutamatergic neurons in the RVM. We analyzed the distribution of these axonal projections in the lumbar spinal cord from Vglut2Cre mice expressing with Cre-dependent mCherry. In contrast to RVM_GABA_ projections, we found almost no RVM_Glu_ processes in LI-II (**Figure 2G, right top**), with mCherry labeling increasing in LIII-V (**Figure 2G, right bottom**). We found dense RVM_Glu_ processes in LX around the central canal (**Figure 2G, bottom left**). Some glutamatergic processes were also present throughout the ventral horn (**Figure 2G, bottom right**). Quantification revealed that fiber densities were significantly greater in LIII-V and LX compared to superficial laminae in the spinal cord (**Figure 2H**). Taken together with the spinal cord projection patterns of RVM_GABA_ neurons, these data indicate that excitatory and inhibitory spinally-projecting RVM neurons may synapse onto different neuronal populations in the dorsal spinal cord to regulate descending pain control. Additionally, the ventral projections of both excitatory and inhibitory neurons suggest an unexamined role for spinally-projecting RVM neurons in modulating motor responses to incoming nociceptive stimuli.

### Anatomical and cell type distribution of the mu opioid receptor in the RVM

The RVM densely expresses mu opioid receptors (MORs), which exert a broadly inhibitory effect on neurons and synapses when activated. Local micro-injection of MOR agonists into the RVM exerts a powerful antinociceptive effect [1,5,19,20,41,45], while activation of putative MOR+ neurons is broadly pro-nociceptive [52]. Thus, we reasoned that loss of MOR signaling in RVM_GABA_ and RVM_Glu_ neurons might recapitulate the effects of chemogenetically activating these neurons after nerve injury by removing the opioid-mediated “brake” on their activity.

Before testing this hypothesis, we directly measured the distribution of MORs across cell types and rostro-caudal levels of the RVM. Previous studies have provided strong evidence that a large proportion of RVM_GABA_ neurons are MOR+, and that a smaller fraction of RVM_Glu_ and serotonergic RVM neurons (RVM_Tph2_) express MORs as well [46,54,59]. However, detailed studies on the spatial distribution of MOR expression throughout the rostro-caudal extent of the RVM are lacking. We sought to quantify the fraction of MOR+ RVM_GABA_, RVM_Glu_, and RVM_Tph2_ neurons and map the distribution of these neurons across the rostro-caudal extent of the RVM.

We performed *in situ* hybridization on RVM sections from C57Bl/6 mice, focusing on sections between 5.6mm-6.4mm posterior to bregma. We labeled all sections for *Oprm1* mRNA, the gene that encodes the MOR. In three separate sub-groups, we co-labeled the sections for *Slc32a1* mRNA (encoding Vgat, a marker of GABAergic neurons; **Figure 3A**), *Slc17a6* mRNA (encoding Vglut2, a marker of glutamatergic neurons; **Figure 3B**), or *Tph2* mRNA (encoding tryptophan hydroxylase 2, a marker of serotonergic neurons; **Figure 3C**). We found that a significantly greater fraction of RVM_GABA_ neurons (approximately 80%) expressed *Oprm1* mRNA compared to RVM_Glu_ neurons, though approximately 40% of RVM_Glu_ neurons were MOR+ (**Figure 3D**). For this analysis, we divided RVM_Tph2_ neurons into midline neurons located in the nucleus raphe magnus (RMg), and lateral neurons located in the lateral nucleus paragigantocellularis (LPGi). We found that approximately 60% of LPGi serotonergic neurons (*Tph2* lateral) were *Oprm1*+, while approximately 40% of RMg serotonergic neurons (*Tph2* medial) expressed MOR. Thus, while some populations (RVM_GABA_ and lateral serotonergic neurons) were much more densely *Oprm1*+, all major neurochemically-defined RVM cell types have a large fraction of MOR-expressing neurons.

**Figure 3.**
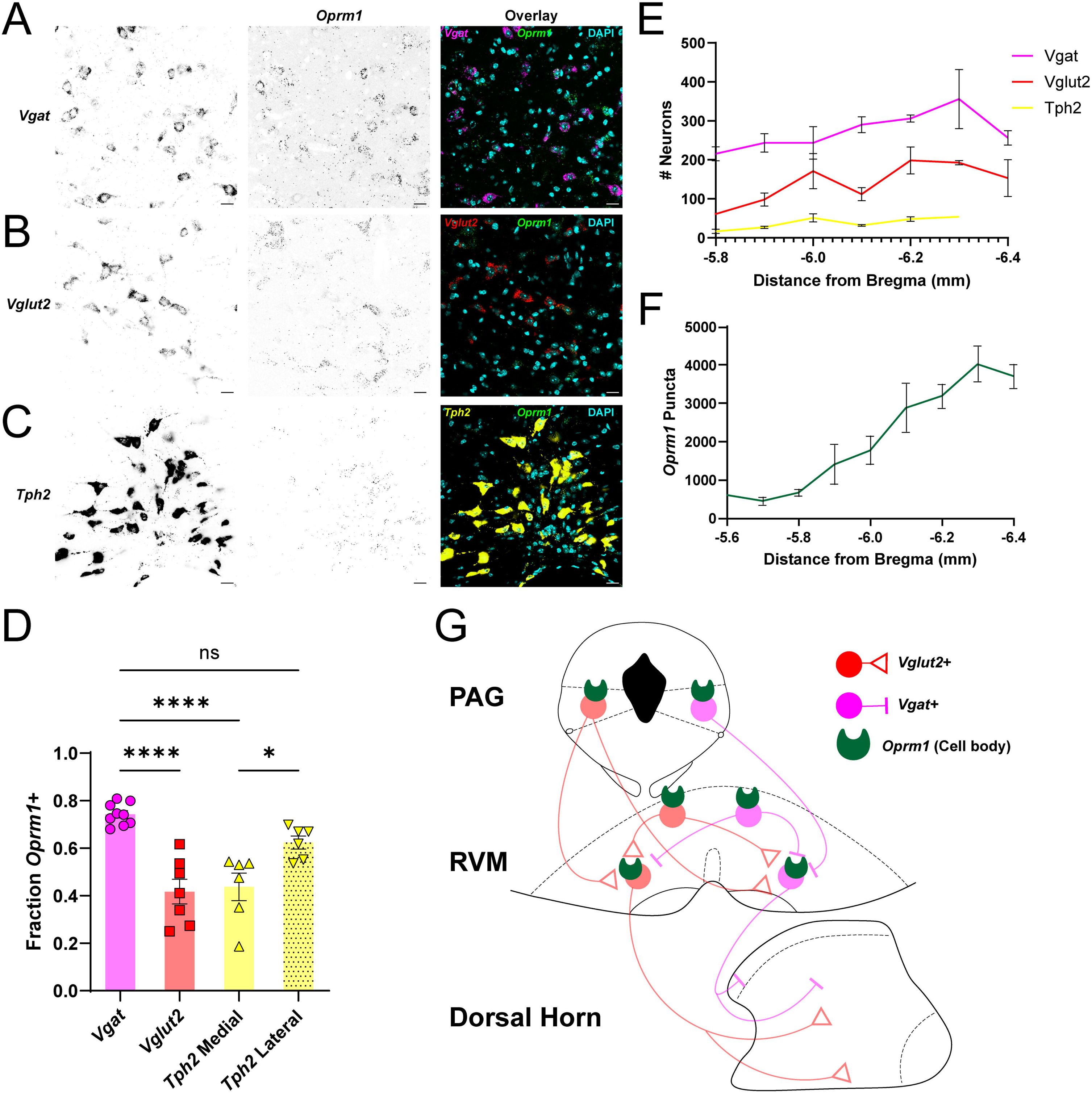
Spatial and cell type distribution of the µ opioid receptor in the RVM. **A,** Representative RVM images showing *Vgat* mRNA expression (left), *Oprm1* mRNA expression (middle), and *Vgat* (magenta)/*Oprm1* (green) overlay (right) (scale bars=20µm). **B,** Representative RVM images showing *Vglut2* mRNA expression (left), *Oprm1* mRNA expression (middle), and *Vglut2* (red)/*Oprm1* (green) overlay (right) (scale bars=20µm). **C,** Representative RVM images showing *Tph2* mRNA expression (left), *Oprm1* mRNA expression (middle), and *Tph2* (yellow)/*Oprm1* (green) overlay (right) (scale bars=20µm). **D,** Quantification of the fraction of *Vgat+*, *Vglut2*+, and *Tph2*+ (medial and lateral) RVM neurons that are *Oprm1+*. One-way ANOVA and post-hoc t-tests with corrections for multiple comparisons; ****p<0.0001, *p<0.05. **E,** Quantification of the number of *Vgat+* (magenta), *Vglut2+* (red), and *Tph2+* (yellow) RVM neurons by distance from Bregma (see Table 3). **F,** Quantification of *Oprm1* mRNA expression by distance from Bregma (see Table 3). **G,** Proposed diagram of RVM *Oprm1* expression and projection patterns of RVM_GABA_ and RVM_Glu_ neurons. Red=Glutamatergic neurons, magenta=GABAergic neurons, green=µ opioid receptors. See **Supplemental Table 3** for detailed statistical analyses.

This *in situ* approach also allowed us to directly map the spatial organization of GABAergic, glutamatergic, serotonergic, and MOR+ neurons across the RVM. We found that, across the rostro-caudal extent of the RVM, the most abundant cell type were RVM_GABA_ neurons, followed by RVM_Glu_ neurons and finally, RVM_Tph2_ neurons (**Figure 3E**). We also analyzed the number of *Oprm1* mRNA puncta across the RVM, and found a strong positive correlation between the distance from bregma and *Oprm1* puncta, indicating greater prevalence of MOR+ neurons in the caudal region of the RVM (**Figure 3F**).

### Cell-type specific knockout of MORs using viral CRISPR/Cas9 gene editing

Having demonstrated that a significant proportion of both RVM_GABA_ and RVM_Glu_ neurons express MOR, we set out to test whether MOR knockout in each of these specific cell types would exacerbate neuropathic pain post-SNI. To achieve this cell type-specific knockout selectively in the RVM, we opted for a CRISPR/Cas9-based approach using a single viral vector to express both the *Oprm1* gRNA and a Cre-dependent fluorophore to visualize edited neurons [49,50].

First, we designed and tested a gRNA against exon 2 of *Oprm1* (**Figure 4A**). We found that our *Oprm1* gRNA candidate (gMOR) produced an insertion or deletion in 80.22% of Neuro2A cells transfected with the guide and spCas9 plasmid (**Figure 4B**). Specifically, gMOR editing resulted in 2.67% insertions, 77.56% deletions, and 1.56% single nucleotide polymorphisms (SNPs) (**Figure 4B**, inset).

**Figure 4.**
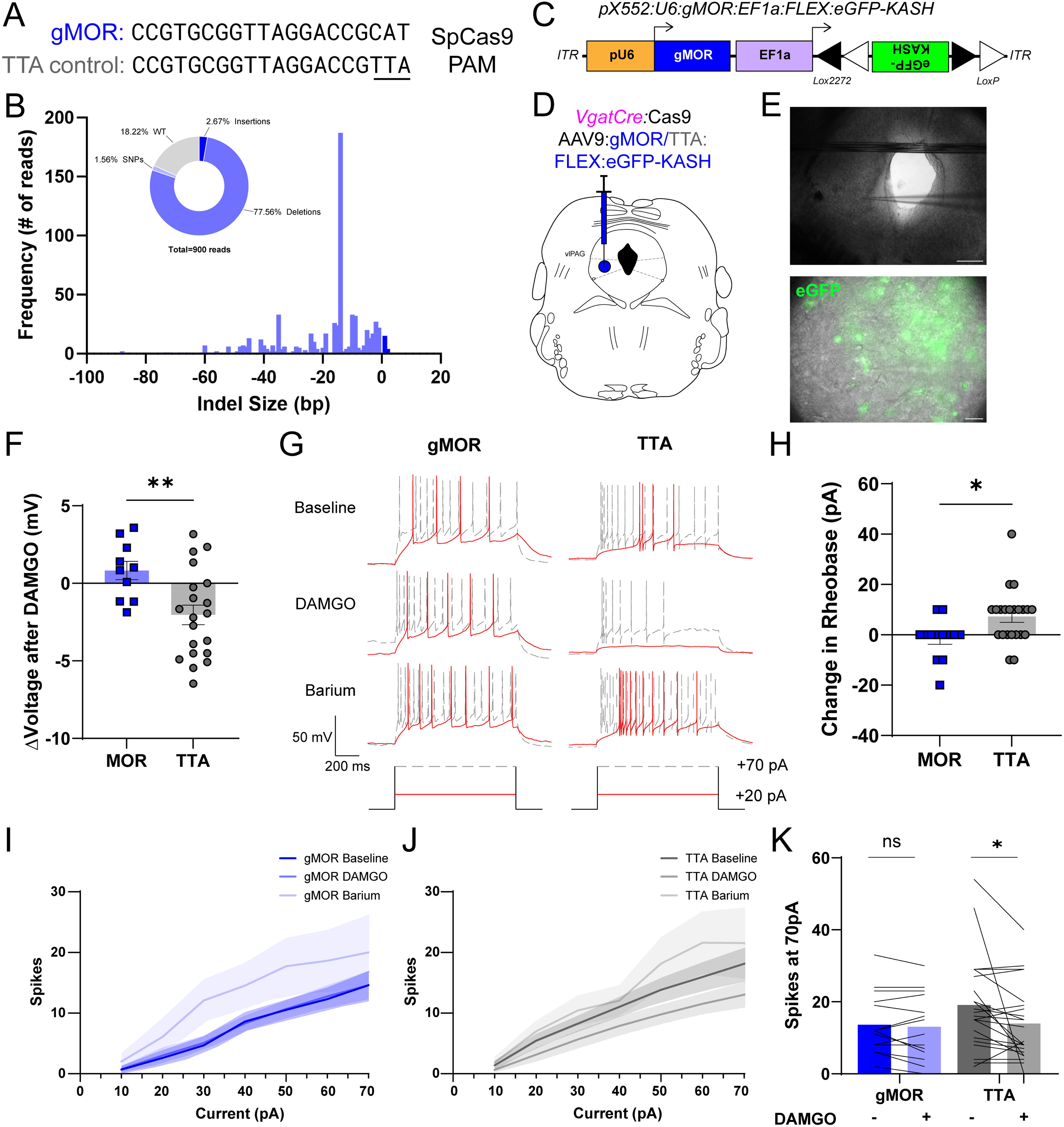
CRISPR/Cas9 gene editing can be used to delete MOR in targeted neuronal populations. **A,** gRNA sequence for CRISPR/Cas9 mediated gene knockout targeted against MOR (gMOR, blue, top) or a control gRNA (TTA control, gray, bottom). **B,** Expression of gMOR *in vitro* efficiently induces indels in the *Oprm1* gene. Main figure demonstrates the frequency of indels in the *Orpm1* gene by size of indel. Inset shows the fraction of total sequencing reads through the *Oprm1* gene that contained insertions (dark blue), deletions (blue), single nucleotide polymorphisms (SNPs; light blue), or wild-type reads (gray). **C,** Diagram of viral vector containing gMOR and Cre-dependent nuclear-targeted eGFP used for electrophysiology and subsequent behavioral experiments. **D,** Diagram of viral injection into the vlPAG of *VgatCre*:Cas9 mice to knock-out MOR in GABAergic vlPAG neurons. **E,** Example images taken during electrophysiology recordings from GABAergic vlPAG neurons. Top, 4X image demonstrating recording site within vlPAG. Scale bar=200µm. Bottom, 40X image demonstrating recording from a transfected GABAergic vlPAG neuron, identified via Cre-dependent eGFP expression. Scale bar=20µm. **F,** Quantification of change in membrane voltage after application of MOR-selective agonist DAMGO in gMOR edited (blue) and TTA control (gray) neurons. 2-tailed t-test, **p<0.01. **G,** Representative traces from gMOR edited (left) and TTA control (right) neurons after injection of a 20pA threshold stimulus (red) and 70pA suprathreshold stimulus (gray) at baseline (top), with DAMGO (middle), and barium (bottom). Horizontal scale bar=200ms, vertical scale bar=50mV. **H,** Quantification of change in rheobase after application of DAMGO in gMOR edited (blue) and TTA control (gray) neurons. 2-tailed Mann-Whitney test, *p<0.05. **I,** Input/output curves for gMOR edited neurons showing average number of spikes/current step at baseline (dark blue), with DAMGO (blue), and barium (light blue). **J,** Input/output curves for TTA control neurons showing average number of spikes/current step at baseline (dark gray), with DAMGO (gray), and barium (light gray). **K,** Quantification of number of spikes at 70pA suprathreshold current injection for gMOR edited (blue) and TTA control (gray) neurons before and after DAMGO application. 2- way ANOVA and post-hoc t-tests with correction for multiple comparisons; *p<0.05. See **Supplemental Table 4** for detailed statistical analyses.

Having confirmed that gMOR can efficiently induce frame-shifting indels *in vitro*, we packaged our candidate gRNA or a control gRNA in an AAV viral vector with Cre-dependent eGFP-KASH (a nuclear targeting sequence) to infect and edit neurons *in vivo* (**Figure 4C**). To test the functional editing capacity of our construct *in vivo*, we targeted GABAergic neurons in the ventrolateral periaqueductal grey (vlPAG), as these neurons densely express MOR and are well-characterized [62,65]. To accomplish this, we crossed Vgat-IRES-Cre mice to mice expressing cre-dependent Cas9 to generate mice that express Cas9 in GABAergic neurons (Vgat-IRES-Cre/H11-LSL-Cas9). We then injected our virus containing either gMOR or TTA control and Cre-dependent eGFP-KASH into the ventrolateral periaqueductal grey (vlPAG) of these mice (**Figure 4D**). This manipulation specifically targets *Oprm1* in GABAergic PAG neurons, which densely express MOR [15,62,65]. After 4 weeks to allow editing and eGFP expression, we performed slice electrophysiology recordings of eGFP+ GABAergic neurons in the vlPAG (**Figure 4E**). Recordings were performed in the presence of synaptic blockers before and after bath application of DAMGO, a MOR-selective agonist, to isolate the effects of MOR knockout in individual neurons.

We first measured the change in membrane voltage of gMOR edited vs. TTA control neurons after bath application of DAMGO. We found that editing *Oprm1* significantly reduced the hyperpolarizing effects of DAMGO on membrane voltage, compared to control neurons (**Figure 4F**). We then measured the change in neuronal excitability before and after DAMGO in response to depolarizing current injections (**Figure 4G**). While neurons from TTA controls exhibited an increased rheobase after DAMGO application, the excitability of Oprm1 edited neurons were not affected (**Figure 4H**), indicating efficient MOR knockout. We also plotted number of APs as a function of current injected for edited and control neurons at baseline, after DAMGO application, and after application of barium, which blocks G protein-coupled inward rectifying potassium (GIRK) channels, a major mechanism by which MORs exert their hyperpolarizing effects at somatodendritic compartments. We found no change in the number of spikes per current step in edited neurons after application of DAMGO (**Figure 4I**), while in control neurons, DAMGO shifted the curve rightward, resulting in fewer APs per current step (**Figure 4J**). We further quantified this change in excitability by comparing the change in number of APs at 70pA for both gMOR edited and TTA control neurons after DAMGO application (**Figure 4K**). We found that there was no change in AP firing in gMOR edited neurons compared to TTA controls. Taken together, these data confirm efficient functional MOR knockout using this viral CRISPR/Cas9 approach.

### MOR knockout in GABAergic RVM neurons exacerbates early pain hypersensitivity after nerve injury

Having validated this cell-specific CRISPR/Cas9 editing of MORs *in vivo*, we returned to the question of whether MOR knockout in specific RVM cell types would affect pain phenotypes at baseline and after nerve injury. Since a high proportion of RVM_GABA_ neurons express MORs, we first assessed the effect of MOR knockout in this population. We injected viral constructs containing gMOR or TTA control gRNA and Cre-dependent eGFP into the RVM of Vgat-IRES-Cre/H11-LSL-Cas9 mice (**Figure 5A,B**). This manipulation targets both local RVM_GABA_ interneurons as well as RVM-to-spinal cord inhibitory projection neurons (**Figure 5C**). Four to 6 weeks following viral injection, we performed baseline behavioral testing to determine the role of MOR signaling on sensory thresholds in the absence of chronic pain (**Figure 5D**). We found no difference between knockout and control groups in the hot plate, von Frey, Hargreaves, acetone, and brush stroke assays (**Figure 5E-H**). Additionally, MOR knockout in RVM_GABA_ neurons had no effect on morphine analgesia in the hot plate assay (**Figure 5E**), suggesting that RVM MORs are minimally engaged in the absence of injury.

**Figure 5.**
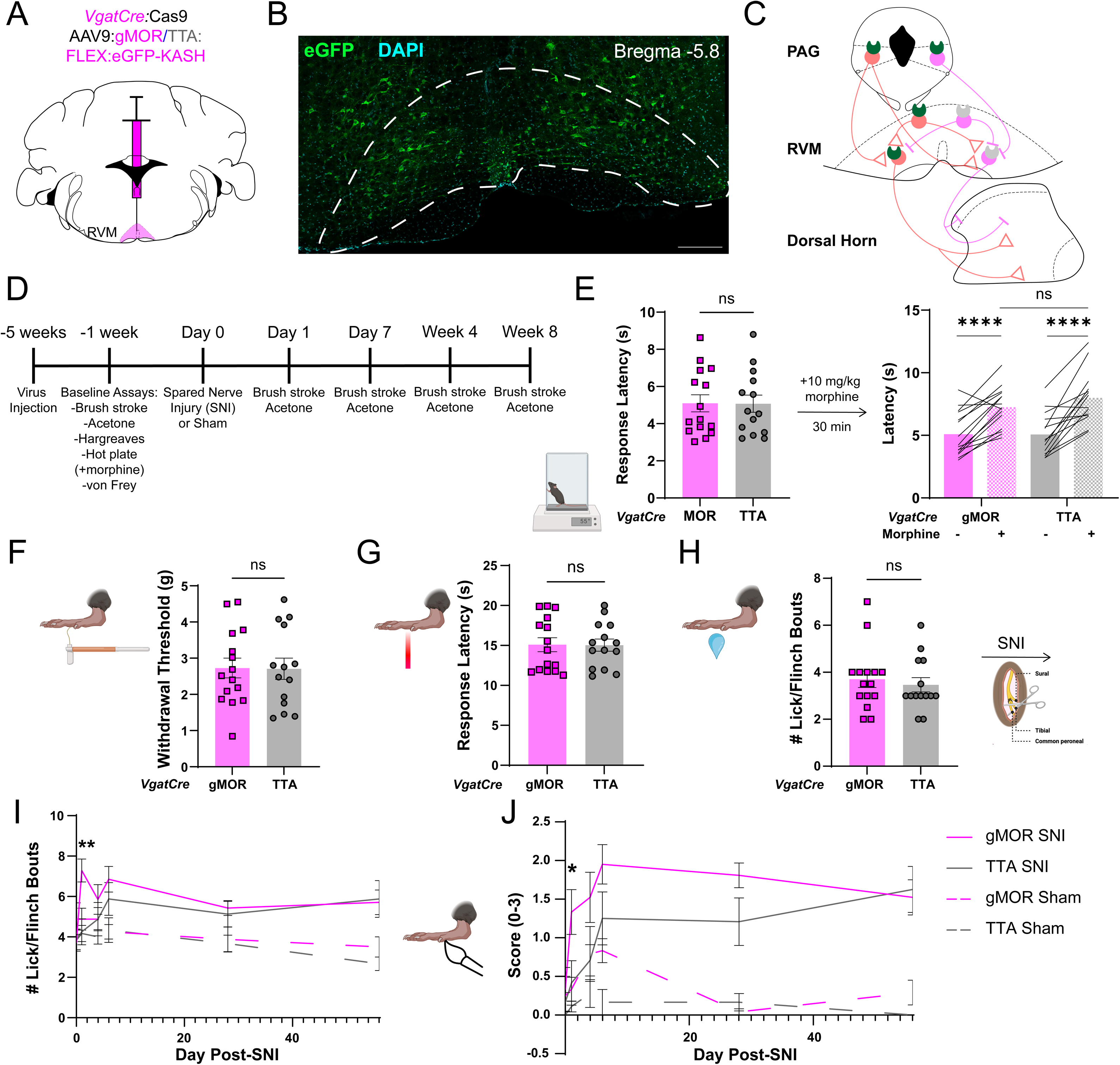
MOR knockout in GABAergic RVM neurons exacerbates pain phenotypes 24h post-nerve injury. **A,** Diagram of injection of active (gMOR, magenta) or control (TTA, gray) virus into the RVM of *VgatCre*:Cas9 mice to knock out MOR in RVM_GABA_ neurons. **B,** Example image of viral expression in RVM_GABA_ neurons at bregma -5.8. Cyan=DAPI, green=Cre-dependent eGFP. Dashed white line indicates RVM. Scale bar=500µm. **C,** Diagram of MOR knockout in RVM_GABA_ interneurons and spinally-projecting neurons. **D,** Timeline of virus injection, behavioral assays, and spared nerve injury (SNI) procedure. **E,** Quantification of hot plate response latency in gMOR edited (magenta) and TTA control (gray) animals before and after administration of 10mg/kg morphine. 2-way ANOVA and post-hoc t-tests with corrections for multiple comparisons; ****p<0.0001. **F,** Quantification of von Frey withdrawal threshold for gMOR edited (magenta) and TTA control (gray) animals. 2-tailed t-test with Welch’s correction. **G,** Quantification of Hargreave’s response latency for gMOR edited (magenta) and TTA control (gray) animals. 2-tailed t-test with Welch’s correction. **H,** Quantification of baseline number of lick/flinch bouts during acetone assay for gMOR edited (magenta) and TTA control (gray) animals. 2-tailed t-test with Welch’s correction. **I,** Quantification of acetone assay licking/flinching bouts for gMOR edited (magenta) and TTA control (gray) mice after SNI (solid line) or sham (dashed line) surgery on days 1, 4 7, 28, and 56 post-SNI. 3-way ANOVA with post-hoc t-tests corrected for multiple comparisons; **p<0.01 for gMOR edited SNI vs. TTA control SNI on Day 1. **J,** Quantification of brush stroke assay score for gMOR edited (magenta) and TTA control (gray) mice after SNI (solid line) or sham (dashed line) surgery on days 1, 4 7, 28, and 56 post-SNI. 3-way ANOVA with post-hoc t-tests corrected for multiple comparisons; *p<0.05 for gMOR edited SNI vs. TTA control SNI on Day 1. See **Supplemental Table 5 and 5.1** for detailed statistical analyses.

To study the role of MOR signaling in this population in chronic neuropathic pain we performed SNI or sham surgery and repeated the acetone and brush stroke assays 1, 4, and 7 days, 4- and 8-weeks post-surgery (**Figure 5D**). In both the acetone and brush stroke assays, MOR knockout in RVM_GABA_ neurons produced an increase in pain responses one day post-SNI when compared to controls (**Figure 5I,J**). This early exacerbation of post-SNI mechanical allodynia and cold hyperalgesia did not lead to persistent increased pain in MOR knockout mice, as there was no difference between knockout and control mice 4 and 7 days post-SNI, or at the 4- and 8-week time points. There were no differences in edited or control mice that received the sham surgery at any time point.

### MOR knockout in glutamatergic RVM neurons exacerbates pain phenotypes after nerve injury

Finally, we examined the effect of *Oprm1* editing specifically in RVM_Glu_ neurons. We injected the gMOR construct or TTA control into the RVM of adult Vglut2-IRES-Cre/H11- LSL-Cas9 mice (**Figure 6A,B**). This manipulation targets both local RVM_Glu_ interneurons as well as RVM-to-spinal cord excitatory projection neurons (**Figure 6C**). Four to 6 weeks following viral injection, we performed baseline sensory testing to determine the role of basal MOR signaling on sensory thresholds and acute reflexive pain responses (**Figure 6D**). Similar to MOR deletion in RVM_GABA_ neurons, we found no difference between knockout and control groups in the hot plate, von Frey, Hargreaves, acetone, and brush stroke assays (**Figure 6E-H**). Additionally, MOR knockout in glutamatergic RVM neurons had no effect on morphine analgesia in the hot plate assay (**Figure 6E**).

**Figure 6.**
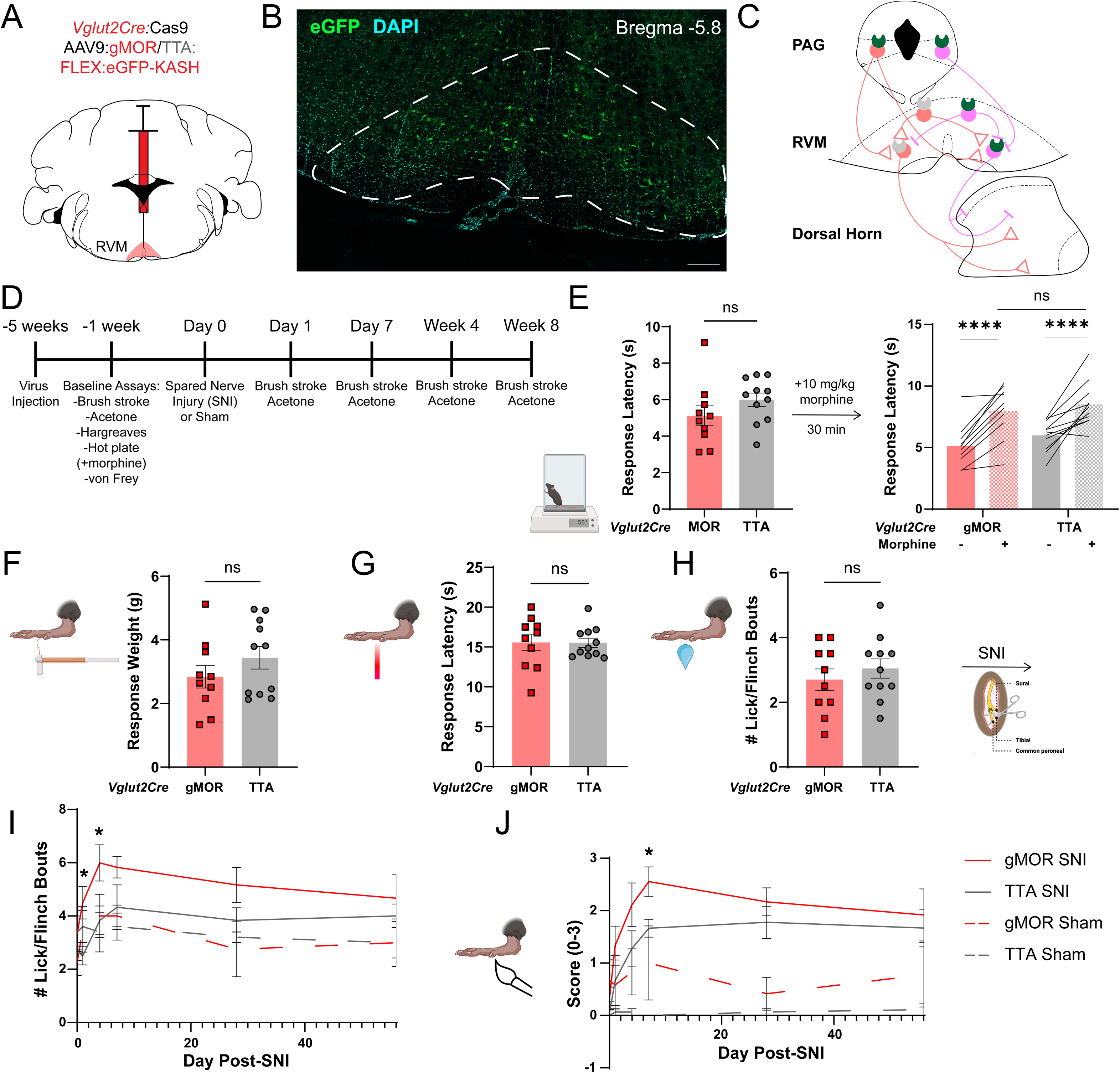
MOR knockout in glutamatergic RVM neurons exacerbates pain phenotypes up to 7 days post-nerve injury. **A,** Diagram of injection of active (gMOR, red) or control (TTA, gray) virus into the RVM of *Vglut2Cre*:Cas9 mice to knock out MOR in RVM_Glu_ neurons. **B,** Example image of viral expression in RVM_Glu_ neurons at bregma -5.8. Cyan=DAPI, green=Cre-dependent eGFP. Dashed white line indicates RVM. Scale bar=500µm. **C,** Diagram of MOR knockout in RVM_Glu_ interneurons and spinally-projecting neurons. **D,** Timeline of virus injection, behavioral assays, and spared nerve injury (SNI) procedure. **E,** Quantification of hot plate response latency in gMOR edited (red) and TTA control (gray) animals before and after administration of 10mg/kg morphine. 2-way ANOVA and post-hoc t-tests with corrections for multiple comparisons; ****p<0.0001. **F,** Quantification of von Frey withdrawal threshold for gMOR edited (red) and TTA control (gray) animals. Nested 2-tailed t-test. **G,** Quantification of Hargreave’s response latency for gMOR edited (red) and TTA control (gray) animals. Nested 2-tailed t-test. **H,** Quantification of baseline number of lick/flinch bouts during acetone assay for gMOR edited (red) and TTA control (gray) animals. 2- tailed t-test with Welch’s correction. **I,** Quantification of acetone assay licking/flinching bouts for gMOR edited (red) and TTA control (gray) mice after SNI (solid line) or sham (dashed line) surgery on days 1, 4 7, 28, and 56 post-SNI. 3-way ANOVA with post-hoc t-tests corrected for multiple comparisons; *p<0.05 for gMOR edited SNI vs. TTA control SNI on Days 1 and 4. **J,** Quantification of brush stroke assay score for gMOR edited (red) and TTA control (gray) mice after SNI (solid line) or sham (dashed line) surgery on days 1, 4 7, 28, and 56 post-SNI. 3-way ANOVA with post-hoc t-tests corrected for multiple comparisons; *p<0.05 for gMOR edited SNI vs. TTA control SNI on Day 7. See **Supplemental Table 6 and 6.1** for detailed statistical analyses.

To study the role of MOR signaling in RVM_Glu_ neurons in chronic neuropathic pain we performed SNI or sham surgery and repeated the acetone and brush stroke assays 1, 4, and 7 days, 4- and 8-weeks post-surgery (**Figure 6D**). In the acetone assay, MOR knockout in RVM_Glu_ neurons produced an increase in pain responses on days 1 and 4 post-SNI compared to control mice that received SNI surgery (**Figure 6I**). In the brush stroke assay, the increased allodynia precipitated by MOR knockout was not apparent until day 7 (**Figure 6J**). In both cases, there was no difference between knockout and control mice at the later 4- and 8-week time points. There were no differences in edited or control mice that received the sham surgery at any time point.

## Discussion

These findings demonstrate the complex role of excitatory and inhibitory RVM neurons in reflexive pain responses at baseline and after neuropathic pain. We show that chemogenetic activation of both RVM_GABA_ and RVM_Glu_ neurons at baseline produces mechanical analgesia; activation of RVM_GABA_ neurons also produces cold analgesia. However, after spared nerve injury (SNI), the analgesic effect of activating RVM_GABA_ neurons is lost. Additionally, we found that RVM_Glu_ neurons became *pro-nociceptive* after SNI. Hypothesizing that RVM MORs may play a role in these post-SNI phenotypes, we demonstrated that the majority of GABAergic RVM neurons and approximately half of glutamatergic and serotonergic RVM neurons express MORs, and that MOR expression is enriched in the more caudal region of the RVM. We demonstrated that cell-specific MOR deletion in the RVM using CRISPR/Cas9 gene editing in both RVM_GABA_ and RVM_Glu_ neurons results in increased mechanical and cold hypersensitivity early (<7 days) post-SNI, with no significant effects at more chronic time points.

These data indicate that excitatory and inhibitory RVM neurons can both contribute to antinociception at baseline, likely through activation or inhibition of distinct groups of spinal cord neurons or presynaptic terminals. Given the almost non-overlapping distributions of GABAergic and glutamatergic RVM projections in the spinal cord, they likely mediate these effects through distinct spinal circuits. Inhibitory GABAergic projections likely inhibit somatic nociceptive signals directly [27,30,53,71] while descending glutamatergic neurons may additionally gate visceral pain [70]. We also observed sparse innervation from the RVM into premotor areas of the spinal cord that may be important for coordinating withdraw or escape behaviors to painful stimuli. Both RVM_GABA_ and RVM_Glu_ neurons also form local connections, allowing for inhibition or activation of local RVM ensembles to produce baseline analgesia. However, our MOR knockout data does not support a significant role for basal MOR signaling in modulating sensory thresholds in excitatory or inhibitory RVM neurons, as cell specific knockout did not affect baseline cold, heat, or mechanical response thresholds.

Deleting MORs specifically in both GABAergic and glutamatergic RVM neurons resulted in enhanced mechanical allodynia and cold hyperalgesia in the acute phase after nerve injury (up to 1 week). However, knockout and control groups experienced similar pain responses at the 4- and 8-week “chronic” time points. These data highlight a critical time window early in the development of chronic neuropathic pain where opioid signaling in the RVM can effectively modulate hypersensitivity and allodynia. The lack of effect of MOR knockout at 4- and 8-weeks post-SNI suggests diminishing efficacy of endogenous opioid modulation in the development of chronic neuropathic pain, potentially mediated by opioid receptor internalization in response to chronically elevated endogenous opioid release. These findings comport with the fact that opioid medications have reduced efficacy in the management of chronic neuropathic pain [4,6,14].

However, RVM MOR knockout did not mirror the effects of chemogenetic activation of RVM_GABA/Glu_ neurons, as we initially hypothesized. MOR knockout resulted in early hyperalgesia after SNI compared to control animals, but the long-term pain outcomes of both groups were similar. Meanwhile, activation of RVM_GABA_ neurons produced no significant effect on pain responses up to 2 weeks post-SNI, and RVM_Glu_ activation exacerbated dynamic allodynia at 2 weeks post-SNI. These findings suggest two conclusions. First, that MOR signaling in the RVM acts in the acute phase of nerve injury to dampen pain signals but has little impact on chronic neuropathic pain. Second, both excitatory and inhibitory MOR+ RVM neurons represent a distinct subset of all RVM neurons, with divergent effects on pain phenotypes[9]. Future studies would be needed to differentiate among possible causes of the decreased efficacy of opioidergic signaling in chronic neuropathic pain, which may include decreased endogenous opioid release, MOR desensitization/internalization, and/or synaptic and circuit restructuring. Additionally, further behavioral studies could more directly test the pro-nociceptive effects of MOR+ RVM neurons after SNI, and compare the connectivity and projection patterns of MOR+ vs. MOR- RVM neurons to better understand how these subpopulations interact to modulate neuropathic pain.

Our findings that MOR knockout in both GABAergic and glutamatergic RVM neurons has a similar effect on neuropathic pain responses suggest that RVM-to-spinal cord projection neurons might be targeting distinct neuronal populations in different laminae of the spinal cord. Indeed, our viral tracing data corroborate this idea, demonstrating an almost non-overlapping spinal projection pattern for inhibitory and excitatory RVM neurons. It is possible that these two RVM neuronal populations exert synergistic effects on nociception by directly inhibiting the central afferents of pain-responsive DRG neurons and/or their postsynaptic targets in the superficial laminae of the spinal cord, and exciting deeper dorsal horn inhibitory interneurons that in turn inhibit projection neurons. This theory could also explain the switch from descending pain inhibition to facilitation we observed with chemogenetic activation of RVM glutamatergic neurons. Changes in chloride gradient in superficial dorsal horn projection neurons have been shown to underly the phenomenon of tactile allodynia in chronic neuropathic pain [26,40,43], resulting in elevated intracellular chloride. Thus, when inhibitory signaling from spinal cord interneurons—driven, in this case, by descending glutamatergic input from the RVM—activates ionotropic GABA_A_ or glycine receptors, this will instead produce a net excitation instead of inhibition. Alternatively, activation of glutamatergic RVM neurons may locally activate serotonergic RVM projection neurons, which have previously been shown to exhibit the same antinociceptive-to-pronociceptive switch after nerve injury [26].

Both GABAergic and glutamatergic RVM neurons send ascending projections to other brain regions involved in pain modulation, such as the PAG [29]. Therefore, the manipulations performed in this study may have had additional effects on other components of the descending pain modulation system that were not explored here. Future studies should examine the role of ascending RVM projections on pain modulation, as this is poorly understood role for the RVM in pain processing. Additionally, the manipulations performed in this paper did not isolate the effects of MOR knockout and chemogenetic activation of local RVM interneurons vs. RVM-spinal cord projection neurons. Several previous studies have investigated the role of spinally- projecting RVM neurons [21,26,28,29,46], and the data in this paper generally corroborate and add to those findings. However, intersectional viral injection techniques to retrogradely label spinally-projecting RVM neurons cannot rule out the role of any local connections or collaterals to other brain regions. The development of new tools to enable selective manipulation of local microcircuits or defined synaptic connections will be important for understanding the key functions of excitatory and inhibitory interneurons in the RVM and other brain regions.

The RVM has been previously shown to mediate changes in pain sensitivity due to stress and other emotional states [10,47,58]. The present study focused on reflexive pain assays as a preliminary exploration of the role of RVM MOR signaling; future studies should examine the effects of RVM MOR knockout on stress induced analgesia and stress induced hyperalgesia, as well as incorporate more affective measures of pain such as conditioned place preference.

This study focused on the role of MORs in RVM neurons that primarily signal using fast neurotransmitters (GABA and glutamate). However, RVM serotonergic neurons also express MORs and are an important component of descending pain modulation [3,19,26,46,67]. Thus, it will be important to determine the role of MOR signaling in this population, as well as dissect the role of “fast” vs. “slow” neurotransmission in the RVM and spinal cord on pain modulation.

Finally, RVM neurons can alternatively be classified by their functional response to pain and opioid analgesia [7,22–24,52]. A more nuanced understanding of these functionally defined populations, including their specific projection targets in the spinal cord, their gene expression profiles, and the role of MOR signaling in these neurons will be important for understanding how the RVM is able to integrate external cues and internal states and modulate nociception. Tools such as fos-TRAP present a powerful mechanism for gaining genetic access to neurons based on their activity in response to specific stimuli [33], and could be used in conjunction with CRISPR/Cas9 gene editing, chemogenetics, or calcium imaging to target functionally-defined neurons in the RVM.

## Conclusion

In this paper, we investigated the role of excitatory and inhibitory RVM neurons, and RVM MOR signaling on baseline reflexive pain responses and in the setting of chronic neuropathic pain. We found that chemogenetic activation of both RVM_GABA_ and RVM_Glu_ neurons at baseline is strongly antinociceptive. Conversely, activation of RVM_GABA_ neurons after SNI has no effect. We also uncover a switch from pain inhibition to pain facilitation in RVM_Glu_ neurons after SNI. We investigated the role of MOR signaling in both of these populations at baseline and after SNI, and found that MORs in both GABAergic and glutamatergic RVM neurons play a key role in antinociception early in the development of neuropathic pain, but not at more chronic time points. Taken together, these findings advance our understanding of the key role the RVM plays as a hub for descending pain modulation, and suggest that interventions which circumvent RVM MOR signaling and selectively inhibit pain facilitating pathways could provide a promising avenue for the treatment of chronic neuropathic pain.

## Supporting information

Statistical Data Table

## Acknowledgements

We would like to thank Dr. Vijay Samineni whose expertise and suggestions helped shape the premise for this study. We would also like to thank all the members of the Copits lab for their helpful discussions and feedback, Rakesh Kumar for assistance and suggestions for SNI and acetone assays, Judy Golden for training and use of the behavior core, the Genome Engineering & Stem Cell Center (GESC@MGI, RRID: SCR_023243) at the Washington University in St. Louis for reagent validation services, and Vera Thornton for consulting on statistical analyses. This work was supported by NIH R01 NS130046 (BAC), NIH R01 DK128475 (VKS) and the McDonnell Center for Systems Neuroscience at WashU Medicine (BAC).

## References

[1] Akaike A, Shibata T, Satoh M, Takagi H. Analgesia induced by microinjection of morphine into, and electrical stimulation of, the nucleus reticularis paragigantocellularis of rat medulla oblongata. Neuropharmacology 1978;17:775–778.

[2] Anderson SD, Basbaum AI, Fields HL. Response of medullary raphe neurons to peripheral stimulation and to systemic opiates. Brain Research 1977;123:363–368.

[3] Andrew H Cooper, Naomi S Hedden, Pranav Prasoon, Yanmei Qi, Taylor BK, Griggs RB. Post-surgical latent pain sensitization is driven by descending serotonergic facilitation and masked by µ-opioid receptor constitutive activity (MOR_CA_) in the rostral ventromedial medulla. The Journal of Neuroscience 2022.

[4] Arnér S, Meyerson BA. Lack of analgesic effect of opioids on neuropathic and idiopathic forms of pain. Pain 1988;33:11–23.

[5] Azami J, Llewelyn MB, Roberts MHT. The contribution of nucleus reticularis paragigantocellularis and nucleus raphe magnus to the analgesia produced by systemically administered morphine, investigated with the microinjection technique. Pain 1982;12:229–246.

[6] Ballantyne JC, Shin NS. Efficacy of Opioids for Chronic Pain: A Review of the Evidence. The Clinical Journal of Pain 2008;24:469–478.

[7] Barbaro NM, Heinricher MM, Fields HL. Putative Nociceptive Modulatory Neurons in the Rostral Ventromedial Medulla of the Rat Display Highly Correlated Firing Patterns. Somatosensory & Motor Research 1989;6:413–425.

[8] Basbaum AI, Fields HL. Endogenous pain control systems: brainstem spinal pathways and endorphin circuitry. Annu Rev Neurosci 1984;7:309–338.

[9] Bekir Nihat Doğrul, Caroline Machado Kopruszinski, Mahdi Dolatyari Eslami, Moe Watanabe, Shizhen Luo, L. H. Moreira de Souza, Robson Lilo Vizin, Xu Yue, Richard D. Palmiter, E. Navratilova, Frank Porreca. Descending facilitation from rostral ventromedial medulla mu opioid receptor-expressing neurons is necessary for maintenance of sensory and affective dimensions of chronic neuropathic pain. Pain 2024.

[10] Butler RK, Finn DP. Stress-induced analgesia. Progress in Neurobiology 2009;88:184–202.

[11] Cai Y-Q, Wang W, Hou Y-Y, Pan ZZ. Optogenetic Activation of Brainstem Serotonergic Neurons Induces Persistent Pain Sensitization. Mol Pain 2014;10:1744–8069-10–70.

[12] Chaplan SR, Bach FW, Pogrel JW, Chung JM, Yaksh TL. Quantitative assessment of tactile allodynia in the rat paw. J Neurosci Methods 1994;53:55–63.

[13] Chiou S-H, Winters IP, Wang J, Naranjo S, Dudgeon C, Tamburini FB, Brady JJ, Yang D, Grüner BM, Chuang C-H, Caswell DR, Zeng H, Chu P, Kim GE, Carpizo DR, Kim SK, Winslow MM. Pancreatic cancer modeling using retrograde viral vector delivery and in vivo CRISPR/Cas9-mediated somatic genome editing. Genes Dev 2015;29:1576–1585.

[14] Chou R, Hartung D, Turner J, Blazina I, Chan B, Levander X, McDonagh M, Selph S, Fu R, Pappas M. Opioid Treatments for Chronic Pain. Agency for Healthcare Research and Quality (AHRQ), 2020 doi:10.23970/AHRQEPCCER229.

[15] Commons KG, Aicher SA, Kow L-M, Pfaff DW. Presynaptic and postsynaptic relations of μ-opioid receptors to γ-aminobutyric acid-immunoreactive and medullary- projecting periaqueductal gray neurons. Journal of Comparative Neurology 2000;419:532–542.

[16] Concordet J-P, Haeussler M. CRISPOR: intuitive guide selection for CRISPR/Cas9 genome editing experiments and screens. Nucleic Acids Research 2018;46:W242–W245.

[17] Dahlhamer J, Lucas J, Zelaya, C, Nahin R, Mackey S, DeBar L, Kerns R, Von Korff M, Porter L, Helmick C. Prevalence of Chronic Pain and High-Impact Chronic Pain Among Adults — United States, 2016. MMWR Morb Mortal Wkly Rep 2018;67:1001–1006.

[18] De Felice M, Sanoja R, Wang R, Vera-Portocarrero L, Oyarzo J, King T, Ossipov MH, Vanderah TW, Lai J, Dussor GO, Fields HL, Price TJ, Porreca F. Engagement of descending inhibition from the rostral ventromedial medulla protects against chronic neuropathic pain. PAIN 2011;152:2701–2709.

[19] Dickenson AH, Oliveras J-L, Besson J-M. Role of the nucleus raphe magnus in opiate analgesia as studied by the microinjection technique in the rat. Brain Research 1979;170:95–111.

[20] Fang FG, Haws CM, Drasner K, Williamson A, Fields HL. Opioid peptides (DAGO-enkephalin, dynorphin A(1–13), BAM 22P) microinjected into the rat brainstem: comparison of their antinociceptive effect and their effect on neuronal firing in the rostral ventromedial medulla. Brain Research 1989;501:116–128.

[21] Fatt MP, Zhang M-D, Kupari J, Altınkök M, Yang Y, Hu Y, Svenningsson P, Ernfors P. Morphine-responsive neurons that regulate mechanical antinociception. Science 2024;385:eado6593.

[22] Fields H, Bry J, Hentall I, Zorman G. The activity of neurons in the rostral medulla of the rat during withdrawal from noxious heat. J Neurosci 1983;3:2545–2552.

[23] Fields HL, Heinricher MM. Anatomy and physiology of a nociceptive modulatory system. Phil Trans R Soc Lond B 1985;308:361–374.

[24] Fields HL, Vanegas H, Hentall ID, Zorman G. Evidence that disinhibition of brain stem neurones contributes to morphine analgesia. Nature 1983;306:684–686.

[25] Follansbee T, Domocos D, Nguyen E, Nguyen A, Bountouvas A, Velasquez L, Iodi Carstens M, Takanami K, Ross SE, Carstens E. Inhibition of itch by neurokinin 1 receptor (Tacr1) -expressing ON cells in the rostral ventromedial medulla in mice. eLife 2022;11:e69626.

[26] Franck Aby, Franck Aby, Lorenzo L-E, Louis-Étienne Lorenzo, Zoé Grivet, Zoé Grivet, Rabia Bouali-Benazzouz, Rabia Bouali-Benazzouz, Hugo Martin, Hugo Martin, Stéphane Valerio, Stéphane Valerio, Sara Whitestone, Sara Whitestone, Dominique Isabel, Dominique Isabel, Idi W, Walid Idi, Bouchatta O, Otmane Bouchatta, De Deurwaerdère P, Philippe De Deurwaerdère, Godin AG, Antoine G. Godin, Herry C, Cyril Herry, Xavier Fioramonti, Xavier Fioramonti, Landry M, Marc Landry, De Koninck Y, Yves De Koninck, Fossat P, Pascal Fossat. Switch of serotonergic descending inhibition into facilitation by a spinal chloride imbalance in neuropathic pain. Science Advances 2022;8.

[27] François A, Low SA, Sypek EI, Christensen AJ, Sotoudeh C, Beier KT, Ramakrishnan C, Ritola KD, Sharif-Naeini R, Deisseroth K, Delp SL, Malenka RC, Luo L, Hantman AW, Scherrer G. A Brainstem-Spinal Cord Inhibitory Circuit for Mechanical Pain Modulation by GABA and Enkephalins. Neuron 2017;93:822–839.e6.

[28] François A, Low SA, Sypek EI, Christensen AJ, Sotoudeh C, Beier KT, Ramakrishnan C, Ritola KD, Sharif-Naeini R, Deisseroth K, Delp SL, Malenka RC, Luo L, Hantman AW, Scherrer G. A Brainstem-Spinal Cord Inhibitory Circuit for Mechanical Pain Modulation by GABA and Enkephalins. Neuron 2017;93:822–839.e6.

[29] Ganley RP, Magalhaes De Sousa M, Ranucci M, Werder K, Öztürk T, Wildner H, Zeilhofer HU. Descending GABAergic Neurons of the RVM That Mediate Widespread Bilateral Antinociception. 2023. doi:10.1101/2023.04.29.538824.

[30] Ganley RP, Sousa M, Ji G, Ranucci M, Beccarini C, Werder K, Pietrafesa F, d’Aquin S, Akyüz T, Hubli M, Schweinhardt P, Neugebauer V, Hoon MA, Wildner H, Zeilhofer HU. Descending inhibitory rostral ventromedial medulla neurons cause widespread antinociception and contribute to the pain-inhibits-pain phenomenon. Nat Commun 2026;17:4765.

[31] Ganley RP, de Sousa MM, Werder K, Öztürk T, Mendes R, Ranucci M, Wildner H, Zeilhofer HU. Targeted anatomical and functional identification of antinociceptive and pronociceptive serotonergic neurons that project to the spinal dorsal horn. eLife 2023;12:e78689.

[32] Gilbert A-K, Franklin KBJ. The role of descending fibers from the rostral ventromedial medulla in opioid analgesia in rats. European Journal of Pharmacology 2002;449:75–84.

[33] Guenthner CJ, Miyamichi K, Yang HH, Heller HC, Luo L. Permanent Genetic Access to Transiently Active Neurons via TRAP: Targeted Recombination in Active Populations. Neuron 2013;78:773–784.

[34] Hargreaves K, Dubner R, Brown F, Flores C, Joris J. A new and sensitive method for measuring thermal nociception in cutaneous hyperalgesia. Pain 1988;32:77–88.

[35] Heinricher MM, Barbaro NM, Fields HL. Putative Nociceptive Modulating Neurons in the Rostral Ventromedial Medulla of the Rat: Firing of On- and Off-Cells Is Related to Nociceptive Responsiveness. Somatosensory & Motor Research 1989;6:427–439.

[36] Heinricher MM, Ingram SL. 5.41 - The Brainstem and Nociceptive Modulation. In: Masland RH, Albright TD, Albright TD, Masland RH, Dallos P, Oertel D, Firestein S, Beauchamp GK, Catherine Bushnell M, Basbaum AI, Kaas JH, Gardner EP, editors. The Senses: A Comprehensive Reference. New York: Academic Press, 2008. pp. 593–626. doi:10.1016/B978-012370880-9.00183-3.

[37] Heinricher MM, Kaplan HJ. GABA-mediated inhibition in rostral ventromedial medulla: role in nociceptive modulation in the lightly anesthetized rat. Pain 1991;47:105–113.

[38] Hunker AC, Soden ME, Krayushkina D, Heymann G, Awatramani R, Zweifel LS. Conditional Single Vector CRISPR/SaCas9 Viruses for Efficient Mutagenesis in the Adult Mouse Nervous System. Cell Reports 2020;30:4303–4316.e6.

[39] Hunker AC, Zweifel LS. Protocol to Design, Clone, and Validate sgRNAs for In Vivo Reverse Genetic Studies. STAR protocols 2020;1:100070.

[40] Jeffrey A. M. Coull, Jeffrey A. M. Coull, Coull JAM, Dominic Boudreau, Boudreau D, Karine Bachand, Bachand K, Steven A. Prescott, Prescott SA, Francine Nault, Nault F, Attila Sı k, Sik A, Paul De Koninck, De Koninck P, Yves De Koninck, De Koninck Y. Trans-synaptic shift in anion gradient in spinal lamina I neurons as a mechanism of neuropathic pain. Nature 2003;424:938–942.

[41] Jensen TS, Yaksh TL. I. Comparison of antinociceptive action of morphine in the periaqueductal gray, medial and paramedial medulla in rat. Brain Research 1986;363:99–113.

[42] Jiao Y, Gao P, Dong L, Ding X, Meng Y, Qian J, Gao T, Wang R, Jiang T, Zhang Y, Kong D, Wu Y, Chen S, Xu S, Tang D, Luo P, Wu M, Meng L, Wen D, Wu C, Zhang G, Shi X, Yu W, Rong W. Molecular identification of bulbospinal ON neurons by GPER, which drives pain and morphine tolerance. J Clin Invest 2023;133. doi:10.1172/JCI154588.

[43] Kaila K, Price TJ, Payne JA, Puskarjov M, Voipio J. Cation-chloride cotransporters in neuronal development, plasticity and disease. Nat Rev Neurosci 2014;15:637–654.

[44] Krashes MJ, Koda S, Ye C, Rogan SC, Adams AC, Cusher DS, Maratos-Flier E, Roth BL, Lowell BB. Rapid, reversible activation of AgRP neurons drives feeding behavior in mice. J Clin Invest 2011;121:1424–1428.

[45] Llewelyn MB, Azami J, Roberts MHT. Brainstem mechanisms of antinociception. Effects of electrical stimulation and injection of morphine into the nucleus raphe magnus. Neuropharmacology 1986;25:727–735.

[46] Marinelli S, Marinelli S, Vaughan CW, Schnell SA, Wessendorf MW, Christie MJ. Rostral ventromedial medulla neurons that project to the spinal cord express multiple opioid receptor phenotypes. The Journal of Neuroscience 2002;22:10847–10855.

[47] Martenson ME, Cetas JS, Heinricher MM. A possible neural basis for stress- induced hyperalgesia. PAIN 2009;142:236–244.

[48] McGaraughty S, Reinis S, Tsoukatos J. Two distinct unit activity responses to morphine in the rostral ventromedial medulla of awake rats. Brain Research 1993;604:331–333.

[49] Moffa J, Kalyanaraman V, Copits B. An Alternative Gene Editing Strategy Using a Single AAV Vector. BIO-PROTOCOL 2025;15. doi:10.21769/BioProtoc.5362.

[50] Moffa JC, Bland IN, Tooley JR, Kalyanaraman V, Heitmeier M, Creed MC, Copits BA. Cell-Specific Single Viral Vector CRISPR/Cas9 Editing and Genetically Encoded Tool Delivery in the Central and Peripheral Nervous Systems. eNeuro 2024;11:ENEURO.0438-23.2024.

[51] Morgan MM, Whittier KL, Hegarty DM, Aicher SA. Periaqueductal gray neurons project to spinally projecting GABAergic neurons in the rostral ventromedial medulla. PAIN 2008;140:376–386.

[52] Neubert MJ, Kincaid W, Heinricher MM. Nociceptive facilitating neurons in the rostral ventromedial medulla. Pain 2004;110:158–165.

[53] Nguyen E, Grajales-Reyes JG, Gereau RW, Ross SE. Cell type-specific dissection of sensory pathways involved in descending modulation. Trends in Neurosciences 2023;46:539–550.

[54] Nguyen E, Holland RA, Ross SE. Cre-based functional profiling of RVM neurons implicates distinct populations in sensory-mediated behaviors. The Journal of Pain 2026;38:105565.

[55] Nguyen E, Smith KM, Cramer N, Holland RA, Bleimeister IH, Flores-Felix K, Silberberg H, Keller A, Le Pichon CE, Ross SE. Medullary kappa-opioid receptor neurons inhibit pain and itch through a descending circuit. Brain 2022;145:2586–2601.

[56] Oliveras J-L, Martin G, Montagne J, Vos B. Single unit activity at ventromedial medulla level in the awake, freely moving rat: effects of noxious heat and light tactile stimuli onto convergent neurons. Brain Research 1990;506:19–30.

[57] Oliveras J-L, Vos B, Martin G, Montagne J. Electrophysiological properties of ventromedial medulla neurons in response to noxious and non-noxious stimuli in the awake, freely moving rat: a single-unit study. Brain Research 1989;486:1–14.

[58] Pagliusi M, Amorim-Marques AP, Lobo MK, Guimarães FS, Lisboa SF, Gomes FV. The rostral ventromedial medulla modulates pain and depression-related behaviors caused by social stress. Pain 2024;165:1814–1823.

[59] Pedersen NP, Vaughan CW, Christie MJ. Opioid receptor modulation of GABAergic and serotonergic spinally projecting neurons of the rostral ventromedial medulla in mice. Journal of Neurophysiology 2011;106:731–740.

[60] Pertovaara A, Wei H, Hämäläinen MM. Lidocaine in the rostroventromedial medulla and the periaqueductal gray attenuates allodynia in neuropathic rats. Neuroscience Letters 1996;218:127–130.

[61] Proudfit HK, Anderson EG. Morphine analgesia: blockade by raphe magnus lesions. Brain Res 1975;98:612–618.

[62] Siuda ER, Copits BA, Schmidt MJ, Baird MA, Al-Hasani R, Planer WJ, Funderburk SC, McCall JG, Gereau RW, Bruchas MR. Spatiotemporal control of opioid signaling and behavior. Neuron 2015;86:923–935.

[63] Stringer C, Wang T, Michaelos M, Pachitariu M. Cellpose: a generalist algorithm for cellular segmentation. Nat Methods 2021;18:100–106.

[64] Vanderah TW, Suenaga NMH, Ossipov MH, Malan TP, Lai J, Porreca F. Tonic Descending Facilitation from the Rostral Ventromedial Medulla Mediates Opioid- Induced Abnormal Pain and Antinociceptive Tolerance. J Neurosci 2001;21:279–286.

[65] Vaughan CW, Bagley EE, Drew GM, Schuller A, Pintar JE, Hack SP, Christie MJ. Cellular actions of opioids on periaqueductal grey neurons from C57B16/J mice and mutant mice lacking MOR-1. British J Pharmacology 2003;139:362–367.

[66] Vong L, Ye C, Yang Z, Choi B, Chua S, Lowell BB. Leptin action on GABAergic neurons prevents obesity and reduces inhibitory tone to POMC neurons. Neuron 2011;71:142–154.

[67] Wei Feng, Wei F, Dubner R, Dubner R, Zou S, Ren K, Bai G, Wei D, Guo W. Molecular depletion of descending serotonin unmasks its novel facilitatory role in the development of persistent pain. The Journal of Neuroscience 2010;30:8624–8636.

[68] Winkler CW, Hermes SM, Chavkin CI, Drake CT, Morrison SF, Aicher SA. Kappa Opioid Receptor (KOR) and GAD67 Immunoreactivity Are Found in off and neutral Cells in the Rostral Ventromedial Medulla. Journal of Neurophysiology 2006;96:3465–3473.

[69] Xu J-J, Gao P, Wu Y, Yin S-Q, Zhu L, Xu S-H, Tang D, Cheung C-W, Jiao Y-F, Yu W-F, Li Y-H, Yang L-Q. G protein-coupled estrogen receptor in the rostral ventromedial medulla contributes to the chronification of postoperative pain. CNS Neuroscience & Therapeutics 2021;27:1313–1326.

[70] Zhang Y, Liu F, Yang L, Hahm HJ, Heitmeier MR, Okuda T, Kawatani M, Harrigan JT, Morrison-Rodriguez EC, Petukhova A, Lu Z, Heitmeier HA, Samineni VK. A Spinal Circuit for Modular Gating of Organ Somatosensation. 2025:2025.08.27.672386. doi:10.1101/2025.08.27.672386.

[71] Zhang Y, Zhao S, Rodriguez E, Takatoh J, Han B-X, Zhou X, Wang F. Identifying local and descending inputs for primary sensory neurons. J Clin Invest 2015;125:3782– 3794.

