## Supplementary material for "The role of mu opioid receptors on excitatory and inhibitory neurons in the rostral ventromedial medulla in neuropathic pain": Statistical Data Table

**Table 1. Statistical data table for Figure 1**

|  | HM3D(Gq) | | | | | | mCherry Control | | | | |
| --- | --- | --- | --- | --- | --- | --- | --- | --- | --- | --- | --- |
| Figure | Saline (mean±SEM) | DCZ (mean±SEM) | | | P value (N) | | Saline (mean±SEM) | | DCZ (mean±SEM) | | P value (N) |
| 1D - # Lick/flinch bouts | 3.1±0.6 | 1.5±0.4 | | | 0.0030** (7) | | 3.0±0.3 | | 2.8±0.4 | | 0.6229 (7) |
| 1E – von Frey withdrawal threshold (g) | 1.03±0.13g | 2.30±0.17g | | | 0.0004*** (7) | | 1.50±0.11g | | 0.98±0.23 | | 0.065 (7) |
| 1F - # lick/flinch bouts post-SNI  Day 1 | 4.3±0.7 | 1.3±0.7 | | | 0.0045** (6) | | 2.6±0.5 | | 2.9±0.4 | | 0.676 (7) |
| Day 7 | 3.5±0.8 | 1.8±0.5 | | | 0.393 (6) | | 3.9±0.6 | | 4.0±0.4 | | 0.850 (7) |
| Day 14 | 4.7±0.7 | 3.2±0.6 | | | 0.067 (6) | | 4.9±0.4 | | 3.1±0.5 | | 0.850 (7) |
| 1G – Brush stroke score post-SNI  Day 1 | 0.03±0.03 | 0.0±0.0 | | | 0.592 (6) | | 0.02±0.02 | | 0.10±0.05 | | 0.829 (7) |
| Day 7 | 1.06±0.10 | 0.94±0.10 | | | 0.918 (6) | | 1.14±0.12 | | 1.00±0.16 | | 0.498 (7) |
| Day 14 | 1.22±0.11 | 0.89±0.22 | | | 0.597 (6) | | 1.14±0.10 | | 1.05±0.05 | | 0.410 (7) |
|  | SNI (mean±SEM) | | | Sham (mean±SEM) | | | | P Value | | | |
| 1I – RVM cFos+ cells/mm^2^ | 146.5±8.7 cells/mm^2^ (n=6 mice) | | | 99.6±21.4 cells/mm^2^ (n=4 mice) | | | | 0.047* | | | |
|  | Lamina I-II (mean±SEM) | | Lamina III-VI (mean±SEM) | | | Lamina X (mean±SEM) | | | | P Value (N) | |
| 1K – RVM_GABA_ projection % Area | 1.65±0.54% | | 0.98±0.22% | | | 1.74±0.41% | | | | 0.1045 (6) | |

**Table 2: Statistical data table for Figure 2**

|  | HM3D(Gq) | | | | | mCherry Control | | | |
| --- | --- | --- | --- | --- | --- | --- | --- | --- | --- |
| Figure | Saline (mean±SEM) | DCZ (mean±SEM) | | P value (N) | | Saline (mean±SEM) | DCZ (mean±SEM) | | P value (N) |
| 2C - # Lick/flinch bouts | 4.4±0.4 | 3.44±0.5 | | 0.1115 (9) | | 4.7±0.5 | 5.3±0.6 | | 0.3840 (8) |
| 2D – von Frey withdrawal threshold (g) | 0.96±0.08g | 2.22±0.12g | | <0.0001**** (9) | | 1.14±0.23g | 0.85±0.16g | | 0.1552 (8) |
| 2E - # lick/flinch bouts post-SNI  Day 1 | 5.3±0.5 | 3.8±1.0 | | 0.1742 (9) | | 4.1±0.7 | 5.4±0.7 | | 0.2500 (8) |
| Day 7 | 4.9±0.7 | 3.9±0.6 | | 0.2891 (9) | | 4.8±0.5 | 3.8±0.3 | | 0.1297 (8) |
| Day 14 | 2.9±0.4 | 3.9±0.3 | | 0.0649 (9) | | 5.6±0.6 | 4.8±0.4 | | 0.2649 (8) |
| 2F – Brush stroke score post-SNI  Day 1 | 1.22±0.30 | 0.63±0.28 | | 0.1640 (9) | | 0.83±0.36 | 0.75±0.34 | | 0.8698 (8) |
| Day 7 | 1.41±0.25 | 2.04±0.34 | | 0.1554 (9) | | 1.04±0.26 | 1.33±0.30 | | 0.4788 (8) |
| Day 14 | 1.19±0.19 | 1.89±0.23 | | 0.0325* (9) | | 1.29±0.17 | 1.13±0.22 | | 0.5579 (8) |
|  | Lamina I-II (mean±SEM) | | Lamina III-VI (mean±SEM) | | Lamina X (mean±SEM) | | | P Value (N) | |
| 2H – RVM_Glu_ projection % Area | 1.06±0.52% | | 3.09±0.64% | | 9.16±2.54% | | | L I-II vs. III-VI: p=0.0045**  L I-II vs. X: p=0.0274*  L III-VI vs. X: p=0.0591 | |

**Table 3: Statistical data table for Figure 3**

| Figure | *Vgat* mean±SEM (N) | | *Vglut2* mean±SEM (N) | | *Tph2* Medial mean±SEM (N) | *Tph2* Lateral mean±SEM (N) | | P Value |
| --- | --- | --- | --- | --- | --- | --- | --- | --- |
| 3D – Fraction of *Oprm1+* neurons | 0.74±0.01 (9 mice) | | 0.42±0.05 (7 mice) | | 0.44±0.06 (6 mice) | 0.62±0.03 (6 mice) | | *Vgat* vs. *Vglut2*: p<0.0001****  *Vgat* vs. *Tph2* medial: p<0.0001****  *Vgat* vs. *Tph2* lateral: p=0.1567  *Vglut2* vs. *Tph2* medial: p=0.9848  *Vglut2* vs. *Tph2* lateral: p=0.0073**  *Tph2* medial vs. *Tph2* lateral: p=0.0218* |
|  | | *Vgat* mean±SEM (N) | | *Vglut2* mean±SEM (N) | | | *Tph2* mean±SEM (N) | |
| 3E - # Neurons  Bregma -5.6 | | 165±0 (1 animal) | |  | | |  | |
| Bregma -5.7 | | 145±11 (2 animals) | | 111±0 (1 animal) | | | 19±0 (1 animal) | |
| Bregma -5.8 | | 216±18 (3 animals) | | 61±0 (1 animal) | | | 17±6 (2 animals) | |
| Bregma -5.9 | | 243±24 (3 animals) | | 98±17 (3 animals) | | | 27±3 (3 animals) | |
| Bregma -6.0 | | 243±42 (3 animals) | | 171±45 (3 animals) | | | 51±11 (3 animals) | |
| Bregma -6.1 | | 290±20 (3 animals) | | 112±17 (2 animals) | | | 32±2 (3 animals) | |
| Bregma -6.2 | | 306±9 (2 animals) | | 199±35 (2 animals) | | | 48±6 (2 animals) | |
| Bregma -6.3 | | 356±76 (3 animals) | | 193±6 (2 animals) | | | 54±0 (1 animal) | |
| Bregma -6.4 | | 257±17 (2 animals) | | 153±47 (3 animals) | | |  | |
|  | | | | *Oprm1* mean±SEM (N) | | | | |
| 3F - # *Oprm1* puncta  Bregma -5.6 | | | | 605±0 (1 animal) | | | | |
| Bregma -5.7 | | | | 451±99 (2 animals) | | | | |
| Bregma -5.8 | | | | 667±95 (3 animals) | | | | |
| Bregma -5.9 | | | | 1408±526 (3 animals) | | | | |
| Bregma -6.0 | | | | 1771±373 (3 animals) | | | | |
| Bregma -6.1 | | | | 2883±635 (3 animals) | | | | |
| Bregma -6.2 | | | | 3197±313 (2 animals) | | | | |
| Bregma -6.3 | | | | 4026±459 (3 animals) | | | | |
| Bregma -6.4 | | | | 3714±320 (2 animals) | | | | |

**Table 4: Statistical data table for Figure 4**

| Figure | | gMOR mean±SEM (N) | | | TTA mean±SEM (N) | | | P value | |
| --- | --- | --- | --- | --- | --- | --- | --- | --- | --- |
| 4F – Change in voltage after DAMGO (mV) | | 0.82±0.59 (10 neurons) | | | -1.72±0.64 (21 neurons) | | | 0.0072** | |
|  | | gMOR median, 25%ile, 75%ile (N) | | | TTA median, 25%ile, 75%ile (N) | | | P value | |
| 4H – Change in rheobase after DAMGO (pA) | | 0, -2.5, 2.5 (10 neurons) | | | 10, 0, 10 (22 neurons) | | | 0.0471* | |
|  | gMOR | | | | | TTA | | | |
| 4I,J - # of spikes per current step | Baseline mean±SEM (N=11) | | DAMGO mean±SEM (N=11) | Barium mean±SEM (N=9) | | Baseline mean±SEM (N=22) | DAMGO mean±SEM (N=22) | | Barium mean±SEM (N=16) |
| 10 pA | 0.727±0.727 | | 0.636±0.636 | 2.222±1.690 | | 1.500±0.850 | 0.682±0.481 | | 2.063±1.109 |
| 20 pA | 2.636±1.527 | | 3.000±1.572 | 6.667±3.790 | | 5.955±1.189 | 3.455±1.230 | | 7.875±1.897 |
| 30 pA | 4.636±1.760 | | 4.909±1.411 | 13.556±4.127 | | 8.636±1.417 | 6.182±1.579 | | 11.313±2.705 |
| 40 pA | 8.909±1.806 | | 8.364±1.454 | 15.889±4.492 | | 11.636±1.738 | 8.591±1.655 | | 13.063±2.762 |
| 50 pA | 11.000±1.945 | | 10.909±1.703 | 19.333±5.612 | | 14.591±2.045 | 10.682±1.624 | | 19.250±4.864 |
| 60 pA | 12.909±2.226 | | 13.182±2.110 | 20.000±6.185 | | 16.864±2.533 | 12.318±1.799 | | 22.125±5.800 |
| 70 pA | 15.455±2.685 | | 15.182±2.713 | 21.444±7.533 | | 19.136±2.760 | 13.955±2.227 | | 21.875±6.585 |
|  | gMOR | | | | | TTA | | | |
|  | Baseline±SEM (N=11) | | DAMGO | P Value | | Baseline | DAMGO | | P Value |
| 4K - # of spikes at 70pA | 15.455±2.685 | | 15.182±2.713 | 0.9215 | | 19.136±2.760 | 13.955±2.227 | | 0.0119* |

**Table 5: Statistical data table for Figure 5E-H**

| Figure | gMOR (N=15 animals) | | | TTA (N=14 animals) | | |
| --- | --- | --- | --- | --- | --- | --- |
|  | Baseline | Morphine | P Value | Baseline | Morphine | P Value |
| Figure 5E - Hot Plate response latency (s) | 5.1±0.5s | 7.2±0.4s | <0.0001**** | 5.1±0.5s | 8.0±0.6s | **TTA baseline vs. morphine:** <0.0001****  **TTA baseline vs. gMOR baseline:** **0.9717**  **TTA morphine vs. gMOR morphine:** 0.2717 |
|  | gMOR (N=15 animals) | | TTA (N=14 animals) | | P Value | |
| Figure 5F – von Frey withdrawal threshold (g) | 2.7±0.3g | | 2.7±0.3g | | 0.9615 | |
| Figure 5G – Hargreaves response latency (s) | 15.1±0.9s | | 15.0±0.8s | | 0.9600 | |
| Figure 5H – Acetone test # lick/flinch bouts | 3.7±0.3 | | 3.5±0.3 | | 0.6136 | |

**Table 5.1: Statistical Data Table for Figure 5I and 5J**

|  | gMOR | | | | TTA | | | |  |
| --- | --- | --- | --- | --- | --- | --- | --- | --- | --- |
|  | SNI (N=7 animals) | P Value (vs. Day 0) | Sham (N=8 animals) | P Value (vs. Day 0) | SNI (N=8 animals) | P Value (vs. Day 0) | Sham (N=6 animals) | P Value (vs. Day 0) | P Value (gMOR SNI vs. TTA SNI) |
| Figure 5I – Acetone test # lick/flinch bouts  Day 0 (Pre-SNI) | 3.7±0.4 | -- | 4.9±0.8 | -- | 3.9±0.5 | -- | 3.8±0.5 | -- | 0.8615 |
| Day 1 Post-SNI | 7.3±0.6 | <0.0001**** | 4.9±0.5 | >0.9999 | 4.3±0.6 | 0.5938 | 4.2±0.4 | 0.9967 | 0.0014** |
| Day 4 | 5.9±0.7 | 0.0056** | 4.9±0.8 | >0.9999 | 4.9±0.6 | 0.1578 | 4.0±0.4 | 0.9999 | 0.2882 |
| Day 6 | 6.9±0.6 | <0.0001**** | 4.3±0.4 | 0.9028 | 5.9±0.8 | 0.0057** | 4.3±0.6 | 0.9788 | 0.2882 |
| Day 28 | 5.4±0.4 | 0.0252* | 3.9±0.6 | 0.5616 | 5.1±0.6 | 0.0787 | 3.7±0.4 | 0.9999 | 0.7419 |
| Day 56 | 5.7±0.6 | 0.0095** | 3.5±0.5 | 0.2161 | 5.9±0.9 | 0.0057* | 2.7±0.3 | 0.5504 | 0.8615 |
|  | SNI (N=7 animals) | P Value (vs. Day 0) | Sham (N=8 animals) | P Value (vs. Day 0) | SNI (N=8 animals) | P Value (vs. Day 0) | Sham (N=6 animals) | P Value (vs. Day 0) | P Value (gMOR SNI vs. TTA SNI) |
| Figure 5J – Brush stroke score  Day 0 (Pre-SNI) | 0.3±0.3 | -- | 0.3±0.3 | -- | 0.2±0.1 | -- | 0.0±0.0 | -- | 0.6033 |
| Day 1 Post-SNI | 1.3±0.3 | 0.1233 | 0.3±0.2 | 0.9975 | 0.4±0.3 | 0.7572 | 0.1±0.1 | 0.6399 | 0.0431* |
| Day 4 | 1.5±0.3 | 0.0695 | 0.8±0.4 | 0.1231 | 0.7±0.2 | 0.2138 | 0.3±0.2 | 0.6623 | 0.0582 |
| Day 6 | 2.0±0.3 | 0.0082** | 0.8±0.2 | 0.3653 | 1.3±0.3 | 0.1528 | 0.2±0.2 | 0.9001 | 0.1263 |
| Day 28 | 1.8±0.2 | 0.0064** | 0.04±0.04 | 0.9042 | 1.2±0.3 | 0.1397 | 0.2±0.1 | 0.6977 | 0.1130 |
| Day 56 | 1.5±0.2 | 0.0185* | 0.3±0.2 | >0.9999 | 1.6±0.3 | 0.0401* | 0.0±0.0 | -- | 0.7921 |

**Table 6: Statistical data table for Figure 6E-H**

| Figure | gMOR (N=10 animals) | | | TTA (N=11 animals) | | |
| --- | --- | --- | --- | --- | --- | --- |
|  | Baseline | Morphine | P Value | Baseline | Morphine | P Value |
| Figure 6E - Hot Plate response latency (s) | 5.1±0.6s | 8.0±0.7s | <0.0001**** | 6.0±0.4s | 8.5±0.6s | **TTA baseline vs. morphine:** <0.0001****  **TTA baseline vs. gMOR baseline:**  0.2546  **TTA morphine vs. gMOR morphine:**  0.4823 |
|  | gMOR (N=10 animals) | | TTA (N=11 animals) | | P Value | |
| Figure 6F – von Frey withdrawal threshold (g) | 2.8±0.4g | | 3.4±0.4g | | 0.2534 | |
| Figure 5G – Hargreaves response latency (s) | 15.6±1.0s | | 15.5±0.6s | | 0.9712 | |
| Figure 5H – Acetone test # lick/flinch bouts | 2.7±0.3 | | 3.0±0.3 | | 0.4503 | |

**Table 6.1: Statistical Data Table for Figure 6I and 6J**

|  | gMOR | | | | TTA | | | |  |
| --- | --- | --- | --- | --- | --- | --- | --- | --- | --- |
|  | SNI (N=6 animals) | P Value (vs. Day 0) | Sham (N=4 animals) | P Value (vs. Day 0) | SNI (N=6 animals) | P Value (vs. Day 0) | Sham (N=5 animals) | P Value (vs. Day 0) | P Value (gMOR SNI vs. TTA SNI) |
| Figure 6I – Acetone test # lick/flinch bouts  Day 0 (Pre-SNI) | 3.3±0.7 | -- | 2.3±0.5 | -- | 2.7±0.3 | -- | 3.4±0.8 | -- | 0.4353 |
| Day 1 Post-SNI | 4.5±0.6 | 0.1410 | 3.5±0.9 | 0.6192 | 2.5±0.3 | 0.8317 | 3.6±0.6 | 0.9994 | 0.0217* |
| Day 4 | 6.0±0.7 | 0.0013** | 4.0±0.8 | 0.5459 | 3.8±0.5 | 0.1410 | 3.4±0.7 | >0.9999 | 0.0132* |
| Day 7 | 5.8±0.4 | 0.0024** | 4.0±0.4 | 0.1532 | 4.3±0.8 | 0.0376* | 3.6±0.5 | 0.9973 | 0.0823 |
| Day 28 | 5.2±0.7 | 0.0228* | 2.8±1.0 | 0.9936 | 3.8±0.5 | 0.1410 | 3.2±0.2 | 0.9999 | 0.1215 |
| Day 56 | 4.7±0.9 | 0.0936 | 3.0±0.6 | 0.8211 | 4.0±0.4 | 0.0936 | 3.0±0.9 | 0.9565 | 0.4353 |
|  | SNI (N=6 animals) | P Value (vs. Day 0) | Sham (N=4 animals) | P Value (vs. Day 0) | SNI (N=6 animals) | P Value (vs. Day 0) | Sham (N=5 animals) | P Value (vs. Day 0) | P Value (gMOR SNI vs. TTA SNI) |
| Figure 6J – Brush stroke score  Day 0 (Pre-SNI) | 0.3±0.2 | -- | 0.7±0.4 | -- | 0.1±0.1 | -- | 0.0±0.0 | -- | 0.5309 |
| Day 1 Post-SNI | 1.3±0.4 | 0.0105* | 0.6±0.5 | 0.9996 | 0.7±0.5 | 0.1106 | 0.1±0.1 | 0.8970 | 0.1356 |
| Day 4 | 2.1±0.4 | <0.0001**** | 0.8±0.4 | 0.5923 | 1.3±0.3 | 0.0021** | 0.1±0.1 | 0.8970 | 0.0635 |
| Day 7 | 2.6±0.3 | <0.0001**** | 1.0±0.7 | 0.8927 | 1.7±0.2 | <0.0001**** | 0.0±0.0 | -- | 0.0482* |
| Day 28 | 2.1±0.3 | <0.0001**** | 0.4±0.3 | 0.6614 | 1.8±0.3 | <0.0001**** | 0.1±0.1 | 0.8970 | 0.3811 |
| Day 56 | 2.1±0.4 | <0.0001**** | 0.6±0.5 | 0.9996 | 1.7±0.2 | <0.0001**** | 0.1±0.1 | 0.6219 | 0.2611 |
